# Structural and functional basis of the universal transcription factor NusG pro-pausing activity in *Mycobacterium tuberculosis*

**DOI:** 10.1101/2022.10.21.513233

**Authors:** Madeleine Delbeau, Expery O. Omollo, Ruby Froom, Steven Koh, Rachel A. Mooney, Mirjana Lilic, Joshua J. Brewer, Jeremy Rock, Seth A. Darst, Elizabeth A. Campbell, Robert Landick

## Abstract

Transcriptional pauses mediate regulation of RNA biogenesis. DNA-encoded pause signals trigger elemental pausing by stabilizing a half-translocated (RNA-not-DNA) state and by promoting RNAP swiveling that other factors can enhance. The universal transcription factor NusG (Spt5 in eukaryotes and archaea) N-terminal domain (NGN) modulates pausing through contacts to RNAP and DNA. Pro-pausing NusGs (e.g., *Bacillus subtilis*) enhance some pauses whereas anti-pausing NusGs (e.g., *Escherichia coli*) suppress some pauses. Little is known about pausing and NusG in the human pathogen *Mycobacterium tuberculosis (Mtb*). Using biochemistry and cryo-electron microscopy, we show that *Mtb*NusG is a pro-pausing NusG that captures paused, swiveled RNAP by contacts to the RNAP protrusion and to a nontemplate strand–DNA wedge inserted between the NGN and the RNAP gate loop. On the other hand, we find that anti-pausing *E. coli* NGN contacts the RNAP gate loop to inhibit swiveling and pausing of *Mtb*RNAP. Using CRISPR-mediated mycobacterial genetics, we show that a pro-pausing NGN is required to support robust mycobacterial growth. Our results define an essential function of NusG in mycobacteria and the structural basis of pro-vs. anti-pausing NusG activity with broad implications for NusG function in all domains of life.

## INTRODUCTION

Regulation of gene expression in all organisms depends on modulating RNA transcript elongation by RNA polymerase (RNAP). To regulate transcript elongation, RNAP evolved to respond to cis-acting pause signals (encoded in the DNA sequence) and to trans-acting regulators that modulate pausing. Similarities in consensus pause signals indicate that pausing has deep evolutionary roots (Gajos et al., 2021; Larson et al., 2014). These pauses play multiple roles in mediating RNA biogenesis. Some pauses enable site-specific regulation of the transcription elongation complex (EC), for example to control premature termination. Many pauses are timing signals that allow nascent RNA folding or interactions of other regulators with the EC (Landick, 2021; Mayer et al., 2017).

At pause sites, variable fractions of ECs can rearrange into elemental paused elongation complexes (ePECs) (Landick, 2006, 2021; Qian et al., 2021; Saba et al., 2019). Structural analyses of *Eco*RNAP and mammalian RNAPII reveal that ePECs form a family of states in which the ~10-bp RNA–DNA hybrid can be pre-translocated or half-translocated (the RNA but not its partner template DNA [t-DNA] nucleotide translocates out of the NTP-binding site) (Guo et al., 2018; Kang et al., 2022; Kang et al., 2018a; Vos et al., 2018). In *Eco*, pause states are also associated with swiveling, an RNAP conformational change in which a set of RNAP structural modules (comprising ~1/3 of RNAP) rotate with respect to the rest of the RNAP. The swiveled RNAP may favor half-translocation and be refractory to nucleotide addition (Kang et al., 2022; Zhu et al., 2022). The ePEC can rearrange further into longer-lived PECs by backtracking or via interactions of nascent RNA structures (pause hairpins, PHs) or trans-acting regulators.

The universal transcriptional regulator NusG can either enhance or suppress pausing (pro-or anti-pausing NusG, Figure 1A) (Kang et al., 2018b; Yakhnin and Babitzke, 2010; Yakhnin et al., 2020b; Zhu et al., 2022). NusG (Spt5 in eukaryotes and archaea) is the only transcription factor found in all three domains of life. NusG, Spt5, and their many paralogs (e.g. RfaH in enteric bacteria) equip ECs for diverse regulatory interactions (Wang and Artsimovitch, 2020). In bacteria, their C-terminal KOW domains contact different transcription regulators such as ribosomes, Rho termination factor, antitermination factors, and RNAP itself (Burmann et al., 2010; Krupp et al., 2019; Lawson et al., 2018; Zhu et al., 2022). All NusG/Spt5 family proteins bind RNAP by contacts of the NusG N-terminal domain (NGN) to antiparallel a-helices on the conserved RNAP clamp domain (clamp helices, CHs). The *Eco*NGN spans the main cleft of RNAP to make additional contacts to the RNAP protrusion, gate loop (GL), or both (Kang et al., 2018b). As a result, the single-stranded nontemplate-strand DNA (nt-DNA) in the transcription bubble threads between the NGN and the clamp.

**Figure 1.**
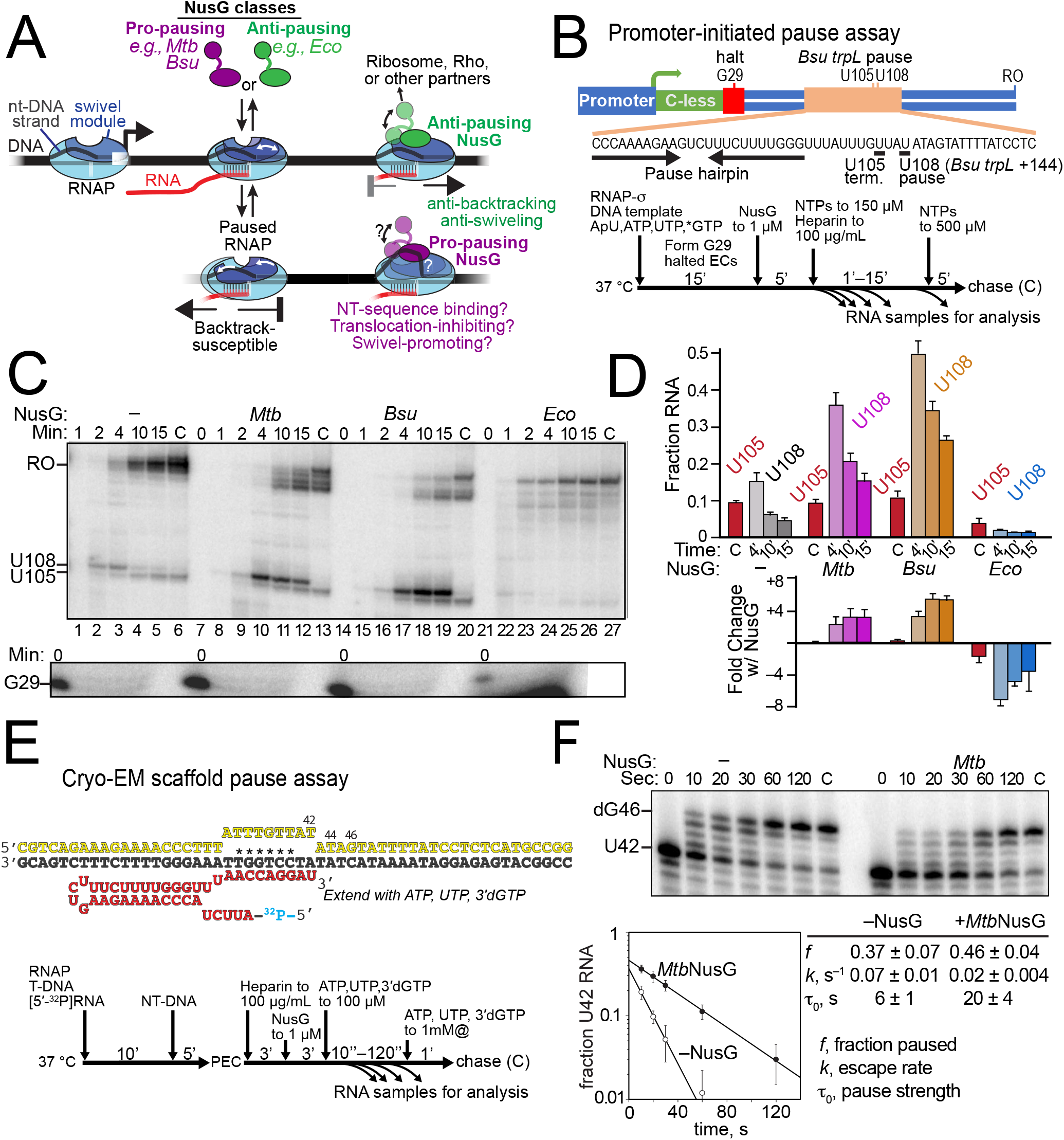
Transcriptional pausing by *Mtb*RNAPand effects of NusG. (A) Mechanism of pausing and effects of pro- and anti-pausing NusGs. (B) Promoter template with *Bsu trpL* pause sequence and site for in vitro transcription assays with *Mtb*RNAP. The template closely resembles one characterized by Yakhnin and Babitzke (2010). The experimental schematic below was followed for data shown in panel C. (C) In vitro pausing by *Mtb*RNAP on promoter-initiated template and effects of pro- and antipausing NusGs. The halted G29 ECs from the same experiment were resolved on a separate gel. The product attributed to U105 RNA did not elongate after addition of 650 μM NTP (C lane) and was therefore attributed to transcriptional termination, whereas the ability of the product U108 to elongate verified assignment as a transcriptional pause. (D) Effects of NusG on pausing and termination. Strength of pausing and termination was assessed by determining the fractions of U105 or U108 RNAs of the sum of U105 plus longer RNAs (e.g., RO RNA). Data are s.d. from 3 experimental replicates. (E) Pause scaffold used for determination of *Mtb*PEC and *Mtb*PEC plus NusG structures (PEC, PEC-MtbNusG, PEC-EcoNusG) and schematic of in vitro pause assays using scaffolds. *, positions at which *Bsu trpL* pause signal was changed to stabilize PEC. (F) Effects of *Mtb*NusG on pausing on cryo-EM pause scaffold. U42 pause RNA was quantified as the fraction of U42 plus longer RNAs (all products that escaped the U42 pause).

NusG was discovered in *Eco* as an anti-pausing factor involved in λN-mediated antitermination (Burova et al., 1995; Li et al., 1992). Alone, anti-pausing *Eco*NusG inhibits backtracking and thus backtrack pausing (Herbert et al., 2010; Kang et al., 2018b; Pasman and von Hippel, 2000; Turtola and Belogurov, 2016). In contrast, *Bacillus subtilis (Bsu), Mycobacterium tuberculosis (Mtb*), and *Thermus thermophilus (Tth*) NusGs exhibit pro-pausing or pro-termination activities (Czyz et al., 2014; Sevostyanova and Artsimovitch, 2010; Yakhnin and Babitzke, 2010; Yakhnin et al., 2020b). Extensive study makes *Bsu*NusG an archetypical pro-pausing NusG (Yakhnin and Babitzke, 2010; Yakhnin et al., 2020b; Yakhnin et al., 2016). Biochemical studies suggest *Bsu*NusG inhibits translocation in PECs via contacts to a TTnTTT motif in the nt-DNA strand. Mammalian Spt5 can exhibit either pro-pausing or anti-pausing activities depending on the action of the P-TEFb kinase (Wada et al., 1998). However, the structural basis of pro-pausing NusG activity remains unknown.

Understanding of the role of NusG in pausing derives primarily from studies of *Eco* and *Bsu*, two distinct but narrow bacterial lineages. Little, if anything, is known about how these mechanisms vary in the huge spectrum of largely unexplored bacterial lineages, notably actinobacteria that includes the major human pathogen *Mtb. Mtb* is the second leading cause of death due to infectious disease worldwide after SARS-CoV-2 (Global_Tuberculosis_Programme, 2021). Here, we show that *Mtb*NusG is pro-pausing with *Mtb*RNAP on a model pause sequence, whereas with the same RNAP and in the same sequence context, *Eco*NusG is anti-pausing. Comparison of *Mtb*NusG to *Eco*NusG allowed us to decipher the structural and mechanistic bases of pro- and anti-pausing and gain insight into fundamental mechanisms of pausing. We determined cryo-EM structures of five *Mtb*RNAP transcription complexes: *i*) an EC; *ii*) a model PH-stabilized PEC; *iii*) the PEC bound by pro-pausing *MtbNusG* (PEC–*Mtb*NusG); *iv*) the PEC bound by anti-pausing *Eco*NusG (PEC–*Eco*NusG); and *v*) the EC bound by *MtbNusG* (EC–*Mtb*NusG). By combining analyses of these structures with biochemistry and CRISPR-mediated genetics, we establish that pro-versus anti-pausing NusG activity results from stabilizing versus suppressing a translocation-inhibited, swiveled PEC state and that the pro-pausing NusG-NGN activity is required for robust cell growth of mycobacteria.

## RESULTS

### Transcriptional pausing by *MtbRNAP* is stimulated by *MtbNusG* at a model pause site

Transcriptional pausing by *Mtb*RNAP is largely uncharacterized, although like nearly all RNAPs tested it recognizes a consensus elemental pause sequence (Larson et al., 2014). Because pausing sequences in *Mtb* have not yet been discovered, we used the well-studied hairpin-stabilized pause signal from the *Bsu trp* operon leader region (*Bsu trpL +144*) to examine the effect of *Mtb*NusG on pausing by *Mtb*RNAP. Extensive analyses by Babitzke and co-workers show *Bsu*RNAP pausing at *trpL* +144, 3 nt after a weak intrinsic terminator at +141, is prolonged by an RNA pause hairpin (PH) that forms upstream from +144U; is prolonged additively by *Bsu*NusG and *Bsu*NusA; and can be conveniently assayed at +108 using shortened templates (Figure 1B) (Yakhnin and Babitzke, 2002, 2010; Yakhnin et al., 2020b). Using the shortened-template approach, we found *Mtb*RNAP also pauses at U108 (*trpL*+144) (Figure 1C, lanes 1–6). Pausing at U108 was enhanced by *Mtb*NusG as well as by *Bsu*NusG (Figures 1C, lanes 8–13, 15–20, and 1D). These pro-pausing NusGs had little effect on intrinsic termination at U105. In contrast, antipausing *Eco*NusG suppressed both U108 pausing and U105 termination (Figures 1C lanes 22–27 and 1D).

To visualize the effect of *Mtb*NusG on the model NusG-stimulated *trpL* PEC, we designed a synthetic RNA–DNA scaffold that included the PH, nt-DNA, active-site and downstream-DNA sequences with six substitutions in the RNA–DNA hybrid to stabilize the PEC without altering the crucial fork-junction sequences (*, Figure 1E). To confirm the synthetic sequence faithfully mimicked NusG-stimulated pausing, we measured the effect of *Mtb*NusG on pause kinetics (Figure 1F). *Mtb*NusG increased τ_0_, a measure of pause strength (Landick et al., 1996), by a factor of ~3.3. We confirmed that the *trpL* PH stimulated pausing by *Mtb*RNAP with or without *Mtb*NusG (Figure S1). We conclude that the synthetic scaffold, encoding the *trpL* pause, permits additive stimulation of pausing by a PH and pro-pausing NusG and is a suitable substrate for cryo-EM analysis of *Mtb*NusG pro-pausing activity.

### The PEC-*Mtb*NusG is half-translocated, swiveled, and stabilized by a nt-DNA wedge

To compare a cryo-EM structure of the PEC-*Mtb*NusG with a canonical EC, we reconstituted *Mtb*RNAP on the synthetic pause scaffold (Figure 1E) and on a scaffold similar to one used previously for a canonical *Eco*RNAP EC (Figure 2A) (Kang et al., 2017). The *MtbEC* scaffold had upstream and downstream DNA extensions relative to the *Eco*EC structure to match the synthetic pause scaffold and programmed G rather than A as the incoming NTP. Incoming G favors a post-translocated EC state (Hein et al., 2011). Both *MtbEC* and PEC-*Mtb*NusG sorted to single conformational states at nominal resolutions of 2.90 Å and 2.94 Å, respectively (Figure S2; Table S1). The *MtbEC* resembled *EcoEC* (Kang et al., 2017) in both the RNAP and post-translocated RNA–DNA conformations, differing notably only in ω interactions with β’, as described previously (Hubin et al., 2017; Lin et al., 2017), and lineage-specific insertions present in the RNAPs (e.g., β’i1 in *Mtb*RNAP clamp; Figure 2A) (Lane and Darst, 2010a). PEC-*Mtb*NusG (Figure 2B) strongly resembled the previously characterized PH-stabilized *Eco hisPEC* (Kang et al., 2018a). Like *Eco hisPEC* and consistent with inability to add NTP rapidly (*i.e*., being paused), PEC-*Mtb*NusG was half-translocated (Figure 2C) and swiveled (Figure 2D). Unlike *his*PEC, which has a 10-bp half-translocated RNA–DNA hybrid and a single nt between the PH and hybrid, the *Mtb*PEC had a 9-bp half-translocated hybrid and at least 3 unresolved nt between the hybrid and the hairpin stem. Like *Eco*RNAP, the *Mtb*RNAP swivel module consists of the shelf, clamp, jaw, dock, and β’Cterm but prominently features rotation of β’i1 in *Mtb*RNAP (Figure 2D), which rotates as part of the clamp, rather than β’SI3 in *Eco*RNAP. (Kang et al., 2018a; Lane and Darst, 2010b)

**Figure 2.**
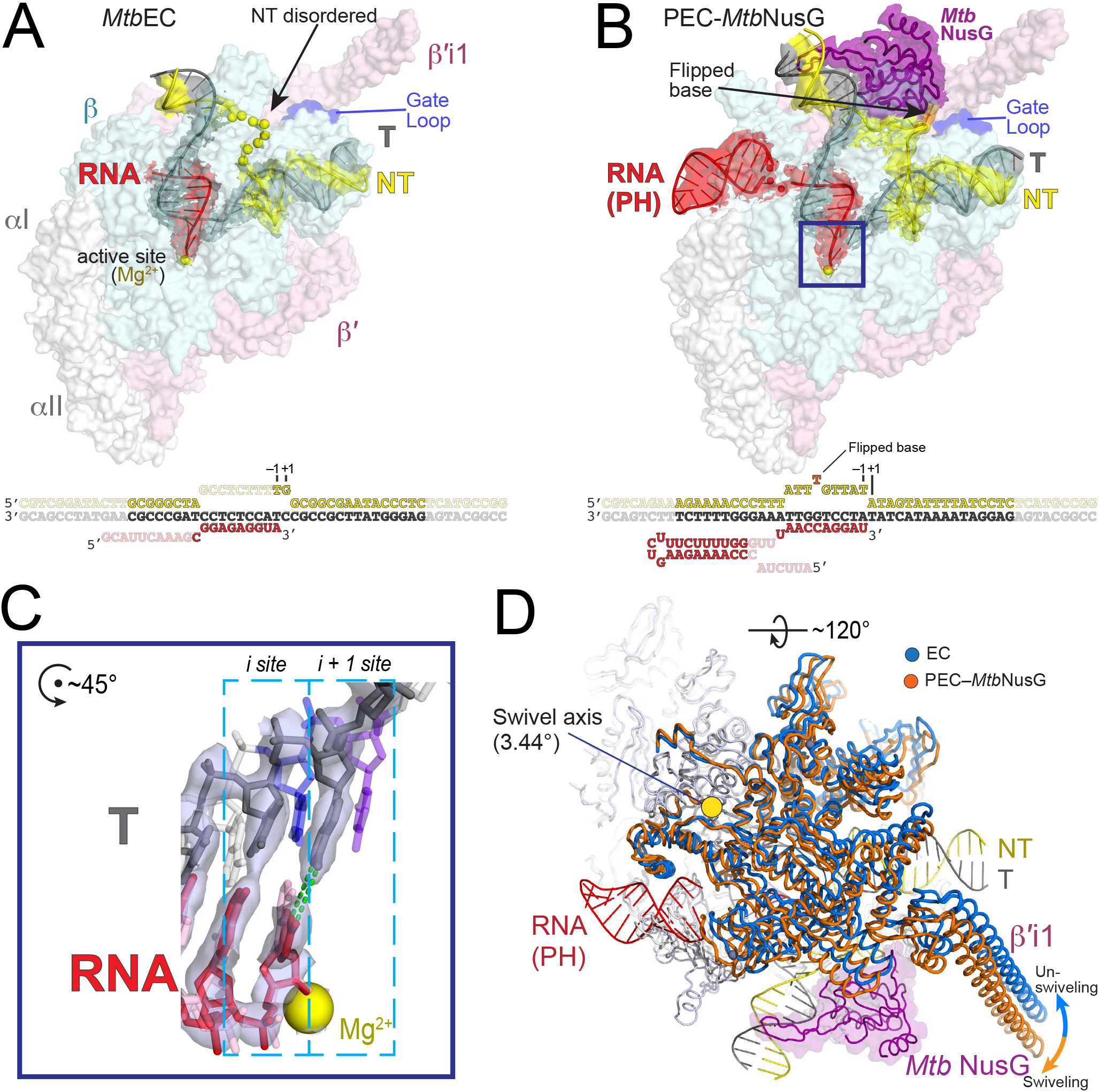
Structural basis of *MtbNusG* pro-pausing activity. (A) Structure of the non-paused *Mtb*RNAP EC. The RNAP is shown as a transparent surface (β’, light pink; β, light cyan; aI, aII, ω, light gray). The transparent local-resolution filtered cryoEM map (Cardone et al., 2013) for the nucleic acids is also shown with the modeled nucleic acids in cartoon (nt-DNA, yellow; t-DNA, dark gray; RNA, red). The EC nucleic acid scaffold is shown in the schematic below (lightly shaded regions have very poor cryo-EM density and are not modeled); the approximate path of the 8-nucleotide single-stranded nt-DNA is denoted with yellow spheres. The complete cryo-EM map is shown in Figure S2D. (B) Structure of paused PEC-*Mtb*NusG. The RNAP is shown as in Figure 2A. The transparent local-resolution filtered cryo-EM map (Cardone et al., 2013) for the nucleic acids and propausing *Mtb*NusG is also shown with the modeled nucleic acids and NusG in cartoon (nt-DNA, yellow; t-DNA, dark gray; RNA, red; pro-pausing *Mtb*nusG, purple). The PEC nucleic acid scaffold is shown in the schematic below (lightly shaded regions have very poor cryo-EM density and are not modeled). The complete cryo-EM map is shown in Figure S2H. (C) Blue-boxed region from panel B zoomed in on the active site. Only nucleic acids and the active site Mg^2+^ are shown for clarity. The half-translocated RNA-DNA hybrid from PEC– *Mtb*NusG is shown (t-DNA, dark gray; RNA, red) along with the cryo-EM density (gray transparent surface). The 3’-nucleotide of the RNA, which is present in the *i site* (posttranslocated), is base paired with the t-DNA nucleotide still in the *i+1 site* (pre-translocated). The post-translocated nucleic acids from EC are shown as reference (t-DNA light gray except t-DNA *i+1 site* is violet and t-DNA *i site* is blue; RNA, light pink). (D) PEC–*Mtb*NusG was aligned to the EC by the RNAP core domain (Boyaci et al., 2018). The structures are shown as backbone worms with the RNAP core domain colored light gray, and the swivel domains colored orange (PEC-*Mtb*NusG) or blue (EC). The PEC-*Mtb*NusG swivel domain (orange) rotates 3.44° clockwise (the view is perpendicular to the rotation axis denoted with the yellow dot) with respect to the EC swivel domain (blue). The nucleic acids and *Mtb*NusG from PEC-*Mtb*NusG are shown (colored according to Figure 2B).

Pro-pausing NusG binds the *Mtb*RNAP CHs similarly to anti-pausing *Eco*NusG binding to *EcoEC* (Kang et al., 2018b; Zhu et al., 2022) and Spt5 to RNAPII or archaeal RNAP (Klein et al., 2011; Martinez-Rucobo et al., 2011), but interacts differently with the nt-DNA and the remainder of RNAP. Swiveling of the swivel module (which includes the CHs) and associated *Mtb*NusG with respect to the rest of the RNAP creates a gap between NusG and the GL into which the single-stranded nt-DNA wedges (Figure 2B). *Mtb*NusG binds the nt-DNA TTATTT motif, with the 3’-T (orange in Figure 2B) flipped into a hydrophobic pocket and stacked on *Mtb*NusG W120, a residue conserved in all NusGs. Unlike the disordered nt-DNA in the *Eco* or *MtbEC* (Figure 2A) and *Eco*NusG EC structures (Kang et al., 2018b; Zhu et al., 2022), the fixed position of the nt-DNA in PEC-*Mtb*NusG both allows its visualization and appears to stabilize the swiveled and thus paused RNAP by wedging between NusG and the GL (Figures 2B, 3D). We conclude that the structural basis of transcriptional pausing by *Mtb*RNAP strongly resembles that determined previously for *Eco*RNAP. Based on the structures of the *MtbEC* and PEC-*Mtb*NusG structures, we conclude that pro-pausing NusG may function, at least in part, by establishing contacts to the nt-DNA that inhibit translocation. Since unswiveling of the RNAP would close the NGN–GL gap that accommodates the wedged nt-DNA (Figures 2B, 3D, and 3E), we also suggest that pro-pausing *Mtb*NusG reinforces swiveling through nt-DNA contacts that stabilize the nt-DNA wedging. Pro-pausing *Mtb*NusG may also reinforce swiveling by stabilizing contacts with other parts of RNAP (discussed in the next section).

**Figure 3.**
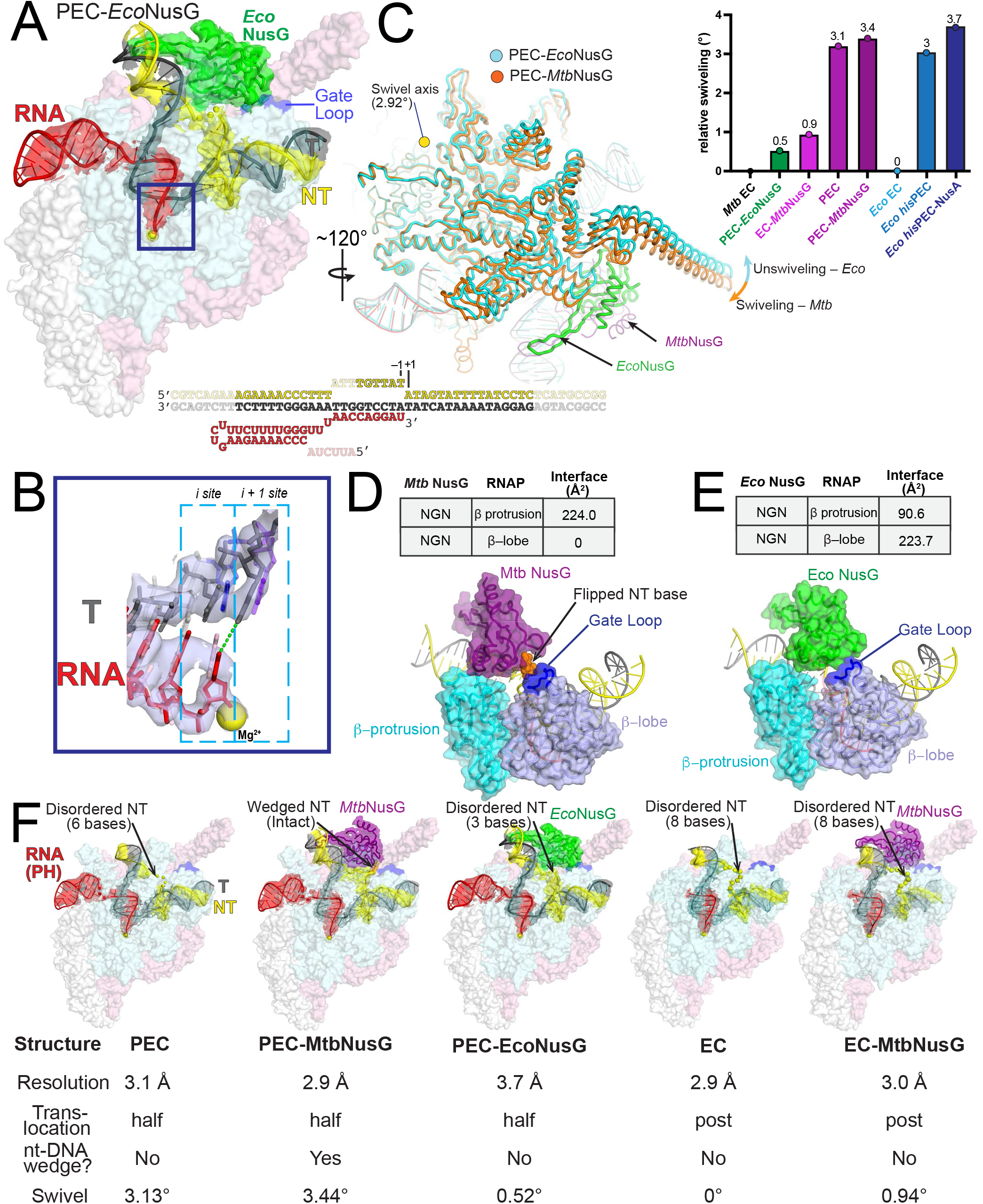
Structural basis of *Eco*NusG anti-pausing activity. (A) Structure of anti-paused PEC-*Eco*NusG. The RNAP is shown as in Figure 2A. The transparent local-resolution filtered cryo-EM map (Cardone et al., 2013) for the nucleic acids and anti-pausing *Eco*NusG is also shown with the modeled nucleic acids and NusG in cartoon (nt-DNA, yellow; t-DNA, dark gray; RNA, red; anti-pausing *Eco*NusG, green). The complete cryoEM map is shown in Figure S3D. (B) Blue-boxed region from panel B zoomed in on the active site. Only nucleic acids and the RNAP active site Mg^2+^ are shown for clarity. The half-translocated RNA-DNA hybrid from PEC–*Eco*NusG is shown (t-DNA, dark gray; RNA, red) along with the cryo-EM density (gray transparent surface). The post-translocated nucleic acids from EC are shown as reference (t-DNA light gray except t-DNA *i+1 site* is violet and t-DNA *i site* is blue; RNA, light pink). (C) PEC–*Eco*NusG was aligned to the PEC-*Mtb*NusG by the RNAP core domain (Boyaci et al., 2018). The structures are shown as backbone worms with the RNAP core domain colored light gray, and the swivel domains colored orange (PEC-*Mtb*NusG) or cyan (PEC-*Eco*NusG). The PEC-*Eco*NusG swivel domain (cyan) rotates 2.92° counter-clockwise (i.e. unswivels; the view is perpendicular to the rotation axis denoted with the yellow dot) with respect to the PEC*-Mtb*NusG swivel domain (orange). The nucleic acids and NusGs from PEC-*Eco*NusG and PEC-*Mtb*NusG are shown. (D, E) Interactions of pro-pausing *Mtb*NusG (D) and anti-pausing *Eco*NusG (E) with the RNAP βprotrusion and βlobe/GL. In both panels, proteins are shown as molecular surfaces (βprotrusion, teal; βlobe, light blue; GL, dark blue). (D) shows the swiveled structure with pro-pausing *Mtb*NusG (purple). *Mtb*NusG forms a substantial interface with the βprotrusion (224 Å^2^ interface area) but does not interact with the βlobe/GL at all. The gap between pro-pausing *Mtb*NusG and the GL accommodates the wedged single-stranded nt-DNA, including the flipped base at the tip of the wedge (shown in orange). (E) shows the unswiveled structure with antipausing *Eco*NusG (green) which forms the most substantial interface with the GL (224 Å^2^). The *Eco*NusG/GL interaction closes the gap accommodating the nt-strand DNA wedge seen in the PEC-*Mtb*NusG structure. (E) All five structures (PEC, PEC-*Mtb*NusG, PEC-*Eco*NusG, EC, EC-*Mtb*NusG) are shown (as in Figures 2A, 2B, and 3A). The nominal resolution, translocation, nt-strand wedging, and swiveling status are denoted below.

### Anti-pausing *Eco*NusG suppresses swiveling

To test our hypothesis that pro-pausing NusG acts at least in part by promoting RNAP swiveling, we determined a cryo-EM structure of anti-pausing *Eco*NusG bound to the *Mtb*PEC (Figures 3A and S3A-D). Like PEC-*Mtb*NusG, the PEC-*Eco*NusG is half-translocated (Figure 3B), but strikingly, *Eco*NusG reverses the swiveling seen in PEC-*Mtb*NusG (Figure 3C). The unswiveled conformation of the PEC-*Eco*NusG eliminates the gap between *Mtb*NusG and the GL seen in the PEC-*Mtb*NusG structure (Figures 3D and 3E), displacing the wedged nt-DNA into a conformation more typical of active ECs.

To ask if NusG directly controls the translocation status of a PEC or EC, we determined additional cryo-EM structures of the PEC alone and the EC plus *Mtb*NusG (Figures 3F, S3E-H, and S4). The PEC was half-translocated and swiveled whereas EC-*Mtb*NusG was posttranslocated and unswiveled (like all ECs so far described); *Mtb*NusG only modestly increased swiveling (Figure 3C; right). These results are consistent with the idea that half-translocation and swiveling are properties of PECs that pro- and anti-pausing NusGs can modulate. Persistence of the half-translocated hybrid in PEC-*Eco*NusG is consistent with the known inability of *Eco*NusG to strongly counteract pause stimulation by PHs (Kang et al., 2018b; Kolb et al., 2014). It is also consistent with the ability of unswiveled but paused RNAPII Spt5-PEC to form a half-translocated hybrid (Vos et al., 2018). Thus, although swiveling may favor half-translocation, it is not a requirement. The at least partial uncoupling of half-translocation and swiveling is consistent with the view that multiple interactions can favor pausing additively in different combinations (Chan et al., 1997; Saba et al., 2019).

The ability of *Mtb*NusG to favor swiveling may also be due to greater interactions between *Mtb*NusG and the protrusion compared to *Eco*NusG, which appears to preferentially interact with the GL (Figures 3D and 3E). Whereas *Eco*NusG has an interface area of ~224 Å^2^ with the βlobe-GL vs ~91 Å^2^ with the protrusion, swiveling eliminates the *Mtb*NusG–βlobe-GL interaction but allows an ~224 Å^2^ interface area with the protrusion (Figures 3D and 3E). These observations suggest that the trade-off between NusG interactions with either the protrusion or the βlobe/GL may be one way that pro-vs. anti-pausing NusGs modulate RNAP swiveling and transcriptional pausing.

### Swiveling may increase pausing by altering the bridge-helix conformation

RNAP swiveling was observed previously in structures of *Eco*RNAP PH-stabilized *his*PECs and Gfh1-containing *TthRNAP* ECs (Guo et al., 2018; Kang et al., 2018a; Sekine et al., 2015). The primary effect of swiveling on *Eco*RNAP pausing arises from inhibition of trigger-loop (TL) folding (Kang et al., 2018a), which is required for efficient phosphoryl-transfer catalysis. Swiveling inhibits TL folding in *Eco*RNAP by shifting a 188-aa insertion in the TL (SI3) into a position that inhibits its movement needed for TL folding (Kang et al., 2018a). Although SI3-like domains are present in the TLs of many bacterial RNAPs (Lane and Darst, 2010a), they are absent in others including *Bsu* and *Mtb*RNAPs that also respond to PHs (Figures 1 and S1) (Yakhnin and Babitzke, 2010; Yakhnin et al., 2008). Swiveling is also associated with non-PH-dependent pausing (Abdelkareem et al., 2019; Guo et al., 2018; Kang et al., 2022; Zhu et al., 2022). These observations raise key questions: *i*) do bacterial RNAPs that lack SI3 still swivel or is swiveling an evolutionary adaptation to the presence of SI3-like domains that allows some bacterial clades to tune their response to pause signals? and *ii*) how do RNAPs lacking SI3 modulate pause lifetimes in response to PHs? Our results answer both these questions.

First, our new structures reveal that *Mtb*RNAP swivels (Figures 2D and 3C). *Mtb* is evolutionarily distant from *Eco* and lacks SI3 (Lane and Darst, 2010a), suggesting that swiveling is a conserved property of bacterial RNAPs that may generally enable PH-stimulation of pausing. Second, *Mtb*RNAP swiveling reveals an attractive explanation for swivel-induced pause enhancement independent of SI3. Rotation of the *Mtb*RNAP swivel module distorts the bridge helix (BH) because the C-terminus of the BH is embedded in the mobile swivel module, whereas the N-terminus of the BH is embedded in the unswiveled body of the RNAP (Figure 4). This BH distortion kinks the middle of the BH and shifts the C-terminal portion of the BH towards the t-DNA, causing steric clash with the optimal position of the post-translocated t-DNA +1 nucleotide (Figure 4). This clash may inhibit translocation of the +1 nucleotide and explain how swiveling increases the energy barrier to conversion of the half-to post-translocated state competent for NTP binding and thus to pause escape.

**Figure 4.**
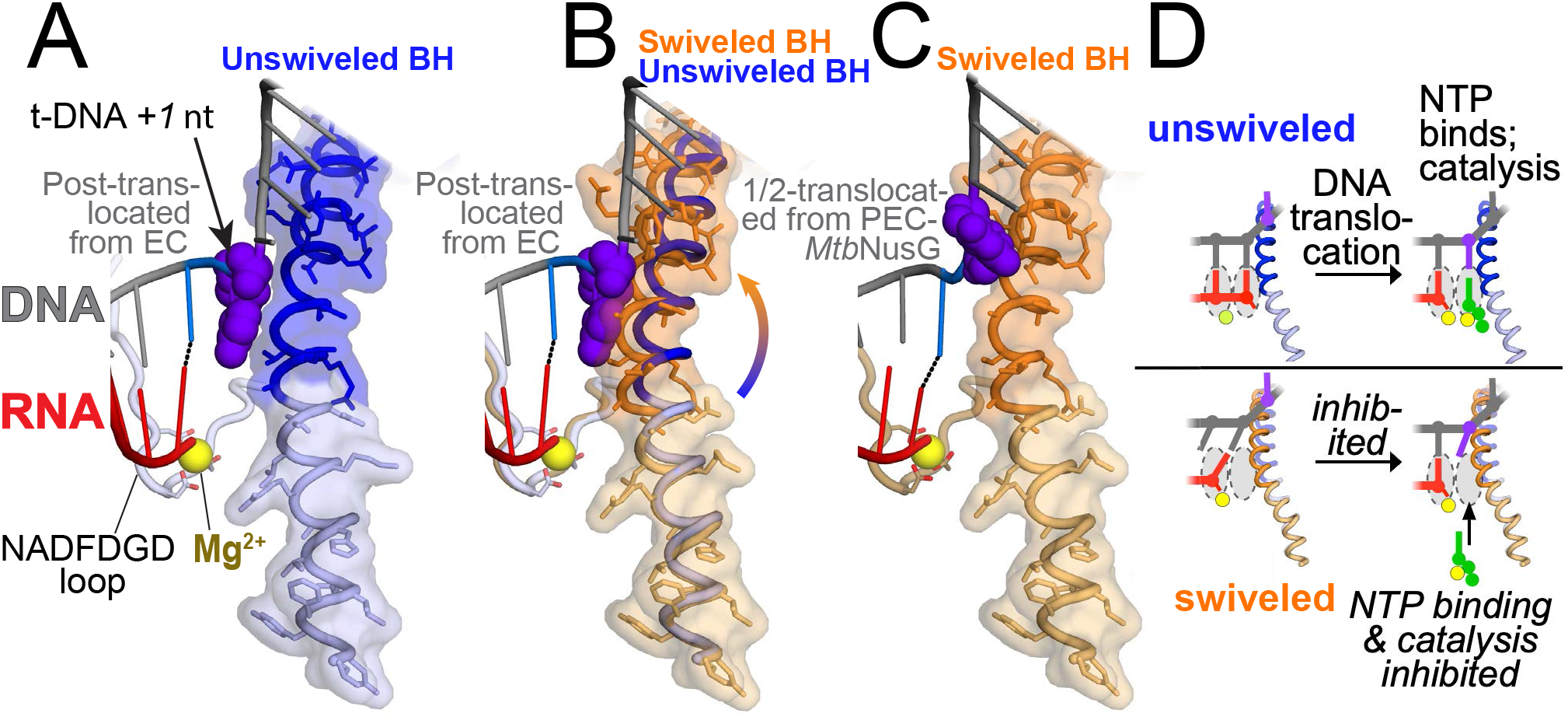
Swiveling, the bridge helix, and pausing. (A) The bridge helix (BH; backbone worm with side chains and transparent molecular surface), the NADFDGD loop (backbone worm with Asp side chains only) and coordinated active site Mg^2+^ (yellow sphere) (Lane and Darst, 2010b), and active site–proximal nucleic acids from the unswiveled EC structure (Figure 2A) are shown. The C-terminal half of the BH (top, blue) is connected to the RNAP swivel domain, while the N-terminal half (bottom, light blue) is connected to the rest of the RNAP. The nucleic acids are post-translocated; the t-DNA nucleotide in the *i+1 site* (which templates the incoming NTP substrate) is shown as purple-blue atomic spheres. (B) The BH and NADFDGD loops from the unswiveled EC (blue backbone worms) and swiveled PEC-*Mtb*NusG (orange backbone worms with transparent orange molecular surface) are superimposed, illustrating the rotation of the BH C-terminal half towards the RNA-DNA hybrid. The nucleic acids are from the unswiveled, post-translocated unswiveled EC to illustrate how the swiveled BH creates a steric clash with the nt-DNA *+1* nucleotide. (C) The BH and NADFDGD loop from the swiveled PEC-*Mtb*NusG are shown with the half-translocated nucleic acids from the same structure, illustrating how the swiveled BH allows for the half-translocated state. (D) Schematic cartoon illustrating how the unswiveled BH (top row, blue) accommodates the pre-translocated (*left*) and post-translocated/NTP-bound states. The swiveled BH (bottom row, orange) accommodates the half-translocated state (*left*). Even if the swiveled RNAP reached the post-translocated state, the nt-DNA *+1* nucleotide would be distorted (with respect to the active site Mg^2+^) to avoid steric clash with the swiveled BH (see panel B), inhibiting NTP binding and/or catalysis.

Additionally, even if DNA translocation is achieved, a swivel-induced BH clash likely inhibits catalysis. Precise positioning of the incoming NTP by the t-DNA +1 nucleotide relative to the catalytic Mg^2+^ ion is essential for rapid catalysis. Mg^2+^ is coordinated by the universally conserved ‘-NADFDGD-’ motif and other conserved active-site elements that are not a part of the swivel module (Figure 4) (Lane and Darst, 2010a). Thus, the kinked BH could preclude proper t-DNA nucleotide positioning and efficient catalysis, thus explaining how swiveling can contribute to transcriptional pausing even in RNAPs that lack SI3. Distortions of the BH and possible effects on translocation and active-site configuration have been noted previously (Bar-Nahum et al., 2005; Gnatt et al., 2001), including in the swiveled, PH-stabilized *his*PEC (Kang et al., 2018a). Our results and analysis clearly reveal why, as postulated (Kang et al., 2022; Zhu et al., 2022), the swiveled state is incompatible with catalysis, and how, in addition to other PEC-specific DNA interactions (Guo et al., 2018; Saba et al., 2019), swiveling prolongs transcriptional pauses.

### Pro-versus anti-pausing NusGs switch conserved side-chain contacts from the GL to nt-DNA and also contact the RNAP protrusion

To ask how pro-vs. anti-pausing activities of NusG depend on NusG sequences, we compared the PEC-*Mtb*NusG and PEC-*Eco*NusG structures and sequences in more detail (Figures 5A, 5B and supplemental data S1). The main contacts between *Mtb*NusG and the wedged nt-DNA are made by W and R residues highly conserved among both pro- and anti-pausing NusGs (Figure 5B). In *Mtb*NusG, W120 forms a surface onto which T6 of the TTATTT motif stacks (Figure 5A). R124 supports W120 through a possible cation-pi interaction with T5 and a guanidinium hydrogen bond to N1 of A3. In the unswiveled PEC-*Eco*NusG structure, the corresponding conserved W and R residues (W80 and R84 of *Eco*NusG) instead contact the GL, along with *Eco*NusG-D77 and H81 (only *Eco*NusG H81 is not conserved between anti- and pro-pausing NusGs; Figure 5B).

**Figure 5.**
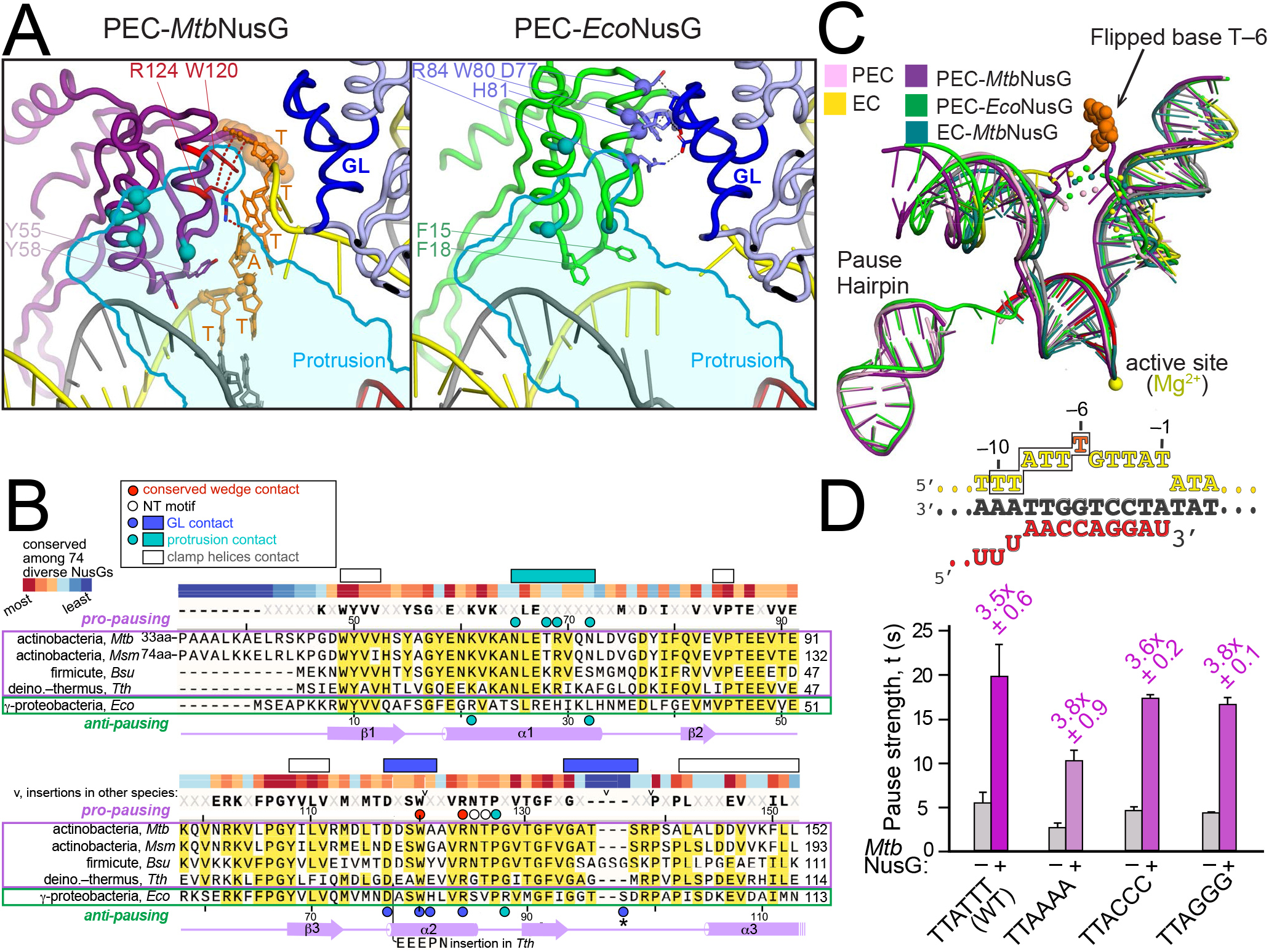
Pro-pausing and anti-pausing NusG interactions with RNAP and nucleic acids. (A) Close up of the swiveled pro-pausing PEC-*Mtb*NusG and unswiveled anti-pausing PEC-*Eco*NusG with residues colored as in the key in panel B. Contacts were identified as described in Methods (supplemental data file 1). (B) Sequence alignment of NusGs with defined pro-pausing and anti-pausing functions. *Mtb, Msm, Bsu and Tth* represent pro-pausing NusGs. The sequence conservation (colored bars and residues in bold) was determined from an alignment of 74 diverse bacterial NusG sequences (supplemental data file 2). *Eco* represents anti-pausing NusGs. Residues of *Mtb*NusG interacting with the *Mtb*RNAP protrusion versus those of *Eco*NusG interacting with the *Mtb*RNAP protrusion and GL are highlighted according to the key. The asterisk indicates a potential H-bond contact that not scored by our computational contact analysis. Residues interacting with the nt-DNA wedged motif from the PEC-*Mtb*NusG structure are also highlighted. (C) All five structures determined here were aligned via the RNAP core module (Boyaci et al., 2018) using the EC (Figure 2A) as the reference. Just the nucleic acids from each structure are shown as cartoons and color-coded as denoted in the legend. The reference EC scaffold is colored as in Figure 2A, and the active site Mg^2+^ is shown as a yellow sphere. The overall configuration of the nucleic acids in each structure is very similar with one exception, segments of the single-stranded nt-strand are disordered except PEC-*Mtb*NusG, where the DNA wedge and flipped –6T are supported by *Mtb*NusG-DNA interactions (Figure 5A). (D) (*top*) Schematic of the section of the pause scaffold shown in (A) highlighting substitutions tested for effects of *Mtb*NusG. (*bottom*) Results of the scaffold-based *in vitro* transcription assay (Figures 1E and 1F) suggest that pro-pausing *Mtb*NusG activity does not require the TTnTTT motif. Data are presented as mean ± SD, n = 3 biologically independent experiments.

It seems unlikely that a single, unconserved His residue in NusG could dictate the switch from pro-to anti-pausing activity. More extensive contacts to the protrusion made by pro-pausing NusG (Figures 3D, 5A, 5B and supplemental data file 1) also likely play a role. Additional residues near the pro-pausing NusG–nt-DNA and anti-pausing NusG–GL interfaces exhibit greater variation than the rest of the NGN and could also support conformations that favor nt-DNA vs. GL contacts. For example, N_125_T_126_ conserved in pro-pausing NusGs vs. S_85_V_86_ in anti-pausing NusGs (*Mtb* vs *Eco* numbering) were proposed to distinguish pro-vs. anti-pausing NusG activities (Yakhnin et al., 2020b). Additionally, *Mtb*NusG Y_55_–Y_58_, which are F_15_–F_18_ in *Eco*NusG, approach the TTnTTT nt-DNA motif and could influence escape from the NusG-stabilized pause (Figure 5A). Our structures suggest that the fundamental basis of pro-pausing activity may depend on multiple differences that include synergy among NusG contacts with the nt-DNA, GL, and protrusion (Figure 5A, supplemental data file 1).

### Pro-pausing activity of MtbNusG may not require the TTnTTT motif

To ask if the pro-pausing activity of MtbNusG depends on the TTnTTT motif as reported for *Bsu*NusG, we examined substitutions in the motif. Somewhat surprisingly, we found that the replacement of the 3’-proximal TTT with AAA, CCC or GGG did not reduce the pro-pausing activity of *Mtb*NusG (AAA did reduce basal pause strength; Figure 5D). The TTnTTT pro-pausing motif for *Bsu*NusG was defined using pause sites that exhibited near-complete dependence on NusG for strong pausing (Yakhnin et al., 2020b), which may reflect interactions important after initial PEC formation (see Discussion).

### Pro-pausing activity of NusG promotes mycobacterial growth

To understand the *in vivo* relevance of NusG pausing activity, we used a CRISPR-interference (CRISPRi) platform previously optimized for mycobacteria (Rock et al., 2017; Wong and Rock, 2021) to transcriptionally silence *nusG* in the model mycobacteria *M. smegmatis (Msm*). In this system, addition of anhydrotetracycline (ATc) induces a nuclease-dead Cas9 (dCas9) and a single-guide RNA (sgRNA) consisting of a dCas9-binding handle and a targeting region complementary to *Msm nusG*, thereby blocking transcription of the endogenous *nusG* locus (Figure S5). By complementing *nusG* knockdown at an ectopic locus, we were able to engineer various “test” *nusG* alleles and examine their effects on cell growth.

We first tested CRISPRi-sensitive versus CRISPRi-resistant *nusG* complementation constructs by comparing the growth of strains in the presence or absence of ATc to induce dCas9 (d’Andrea et al., 2022) (Figure S5). The CRISPRi-resistant *nusG* contained silent mutations to abrogate dCas9 binding without changing the amino acid sequence. Induction of the sgRNA and dCas9 prevented growth of cells expressing CRISPRi-sensitive but not CRISPRi-resistant *nusG* (Figure 6A), consistent with the behavior of this sgRNA in a genome-wide CRISPRi screen in *Msm* (Bosch et al., 2021). We found that *nusG* knockdown could only be complemented by providing the entire predicted transcriptional unit flanking *nusG*, which includes two upstream and two downstream genes (Figures S5A and B). All five genes in the *nusG* transcriptional unit are predicted to be required for mycobacterial viability (Dragset et al., 2019); it is thus likely that *nusG* knockdown induces both upstream and downstream polar effects (Bosch et al., 2021; Peters et al., 2016). Hence, for the purposes of this study, we refer to the genomic region included in the complementation construct as the “*nusG* operon”.

**Figure 6.**
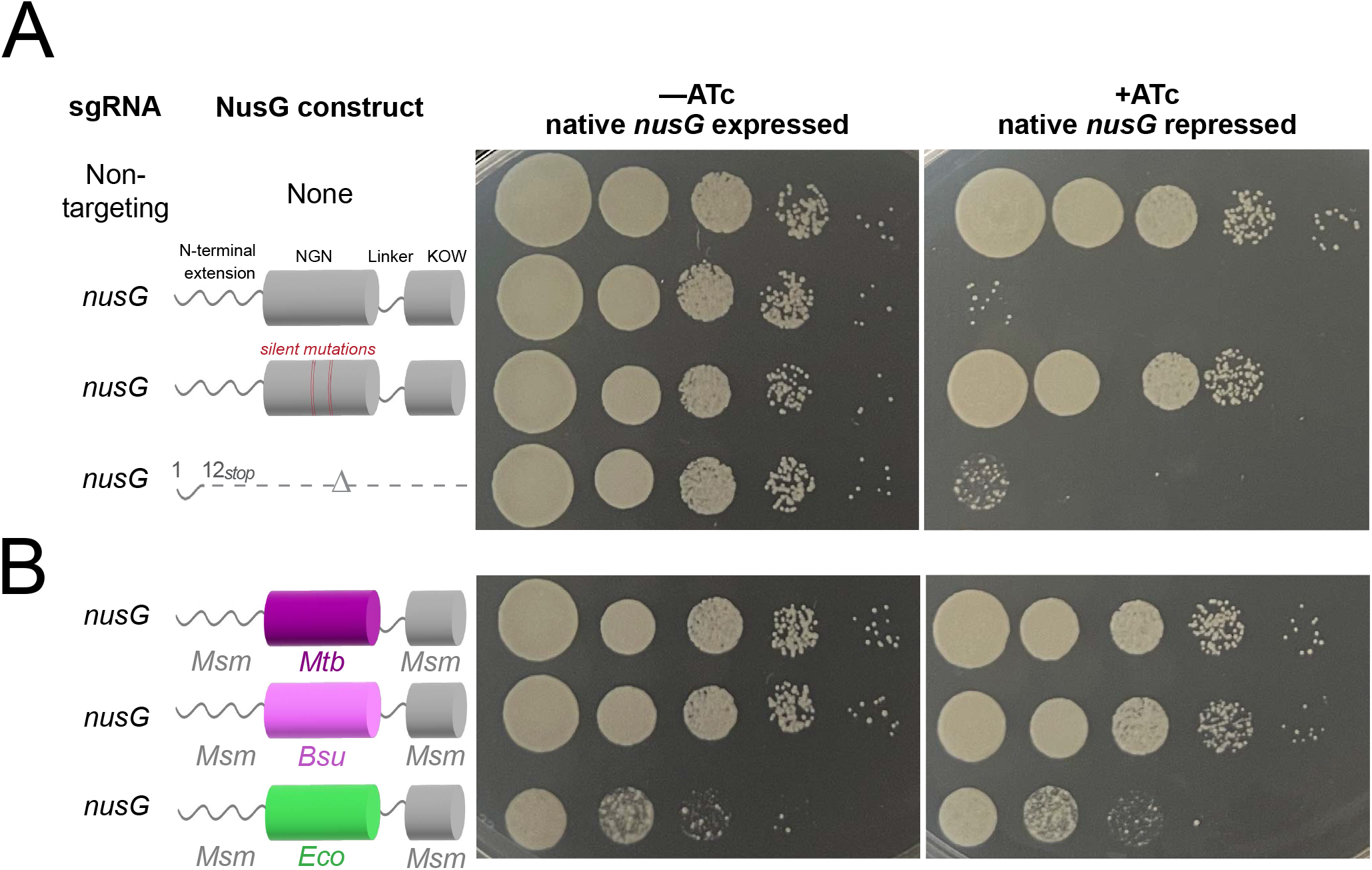
NusG effects on *M. smegmatis (Msm*) fitness. On the left, schematics illustrate the “test” NusG constructs used to complement endogenous nusG knockdowns. On the right, ten-fold serial dilutions of *Msm* strains containing CRISPRi knockdown plasmids and ectopic complementation plasmids were plated in the presence or absence of CRISPRi inducer anhydrotetracycline (ATc) and imaged after two days. (A) *nusG* is an essential gene in *Msm*. Introduction of a premature stop codon in the CRISPRi-resistant *nusG* complementation allele (denoted by “silent mutations”) interferes with *Msm* growth in the context of native *nusG* knockdown. (B) (B) Pausing activity of NusG NGN domains correlates with effects on *Msm* fitness. Pro-pausing NGN fusion constructs (*Mtb, Bsu*) can sustain growth comparable to wild-type *Msm* NusG, while the *Eco* anti-pausing domain interferes with growth.

To isolate the *nusG-specific* contribution to *Msm* growth, we complemented *nusG* operon knockdown with a construct containing a CRISPRi-resistant *nusG* allele with a premature stop codon (D12*) (Chen et al., 2012). This strain shows a strong growth defect upon endogenous *nusG* depletion, consistent with the predicted essentiality of *nusG* (Figure 6A). Immunoblot confirmed that full-length NusG protein levels were depleted in this strain after 12 hours (Figure S6C).

Since the absence of NusG caused a growth defect, this genetic system allowed us to specifically assess how the NGN (necessary and sufficient to modulate pausing activity *in vitro*) affects mycobacterial growth. To this end, we designed complementation constructs where NGNs with known pausing activities from different bacteria were swapped into an otherwise native *Msm*NusG in a way that preserved the predicted NGN structure (Figure 6B and S5D). By preserving the *Msm* N-terminal extension (a disordered Actinobacteria-specific appendage of unknown function), linker and KOW domain (in other species, responsible for bridging RNAP and interactors such as Rho and the ribosome), we sought to measure the fitness effects of pro- vs. anti-pausing NGNs without disrupting other cellular roles of *Msm*NusG.

We found that *nusG* fusion constructs containing pro-pausing NGNs (*Mtb* and *Bsu*) sustain *Msm* growth at levels comparable to wildtype *Msm*NusG (Figure 6B), with or without CRISPRi induction. In striking contrast, expression of the *nusG* fusion construct containing an antipausing *Eco*NGN dramatically decreased cell growth. Notably, this occurred even in the absence of endogenous *nusG* knockdown, consistent with the stability of the *Eco* NusG-*Mtb* RNAP complex. Immunoblots confirmed that all fusion constructs were expressed at similar levels to *Msm*NusG (Figure S5E). Overall, our findings suggest that NusG-stimulated pausing is required for optimal mycobacterial fitness.

## DISCUSSION

We sought to understand *Mtb*RNAP pausing, the mechanistic basis of pro-versus anti-pausing NusG, and the role of the presumptive pro-pausing *Mtb*NusG in mycobacterial growth. Our results, combined with earlier studies, established that *Mtb*RNAP structurally and functionally responds to pause signals and PHs similarly to *Eco*RNAP. Pause sequences dictate an elemental pause state in which translocation is inhibited (here in the half-translocated state) and the PEC is swiveled, likely due to the formation of the PH in the RNAP RNA exit channel (Kang et al., 2018a). We establish that *Mtb*NusG is a pro-pausing NusG that aids RNAP swiveling through contacts to the nt-DNA and the RNAP protrusion, in contrast to anti-pausing *Eco*NusG, which forms contacts to the RNAP βlobe-GL. The pro-pausing activity of *Mtb*NusG is necessary for robust mycobacterial growth based on the ability of pro-pausing but not anti-pausing NGNs inserted in full-length mycobacterial NusG to promote cell growth. These findings provide multiple mechanistic insights and define directions for future research.

### *Mtb*RNAP uses a conserved pause mechanism in which swiveling distorts the BH

A fundamental mechanism of transcriptional pausing in bacteria that involves consensussequence inhibition of DNA translocation and rotation of a subset of RNAP modules has emerged from recent studies of *Eco*RNAP (Abdelkareem et al., 2019; Guo et al., 2018; Kang et al., 2022; Kang et al., 2018a; Kang et al., 2019; Landick, 2021; Qian et al., 2021; Saba et al., 2019; Zhu et al., 2022). Despite evidence that the consensus pause sequence is recognized by many RNAPs (Gajos et al., 2021; Larson et al., 2014), it has remained unclear if the fundamental mechanism is as widespread as the consensus sequence and how swiveling could contribute to pausing in bacteria lacking the SI3 TL insertion. Our finding that *Mtb*RNAP both swivels and inhibits DNA translocation like *Eco*RNAP establishes that these mechanistic features are broadly distributed among bacteria. PHs also stabilize the swiveled *Mtb*PEC similarly to the *Eco his* PH through interactions of the PH with the RNA exit channel and clamp (Figures 2 and 3). By examining the consequences of swiveling in *Mtb*RNAP, which lacks the SI3 TL insertion crucial for PH effects in *Eco*RNAP, we identified a second way swiveling affects the enzyme. Swiveling shifts the portion of the BH anchored in the clamp domain, causing the BH to kink at the connection to the immobile part of the BH. Highly conserved BH Thr–Ala residues near the kink, which normally contact the t-DNA base to position the incoming NTP for catalysis, instead occlude this portion of the active site. This swivel-induced BH occlusion inactivates RNAP in two ways (Figure 4). First, translocation of the next t-DNA base is likely inhibited because the BH Thr-Ala residues sterically preclude translocation. This effect explains how swiveling would stabilize the half-translocated RNA–DNA hybrid, a characteristic of some PECs. Second, even if translocation is achieved, the t-DNA base would be forced into a misaligned position where it would position the incoming NTP suboptimally for catalysis. These insights suggest that swiveling, in addition to inhibiting TL folding when a TL insertion is present, will prove to be a broadly distributed component of pause mechanisms among bacteria, including those lacking a TL insertion, with a common BH-mediated effect on the RNAP active site.

The five cryo-EM structures we determined [Figures 3C and 7); *i*) EC, *ii*) EC-*Mtb*NusG, *iii*) PEC, *iv*) PEC-*Mtb*NusG, *v*) PEC-*Eco*NusG] combined with other recent results (Kang et al., 2022; Zhu et al., 2022) lead us to make the following conjectures:

i. All the ECs, with or without *Mtb*NusG, are post-translocated, while all the PECs, with or without NusGs, are half-translocated. We conclude that sequence characteristics of the upstream- and downstream-fork junctions of the RNA-DNA hybrid induce the half-translocated state (Larson et al., 2014; Saba et al., 2019).
ii. All ECs or PECs explore unswiveled and swiveled states but with different population distributions. ECs spend most of their time unswiveled, while PECs can be biased towards swiveled states to various degrees (Figure 3C). The presence of the PH, such as in the PEC, biases the conformational landscape towards swiveling (and thus towards pausing).

**Figure 7.**
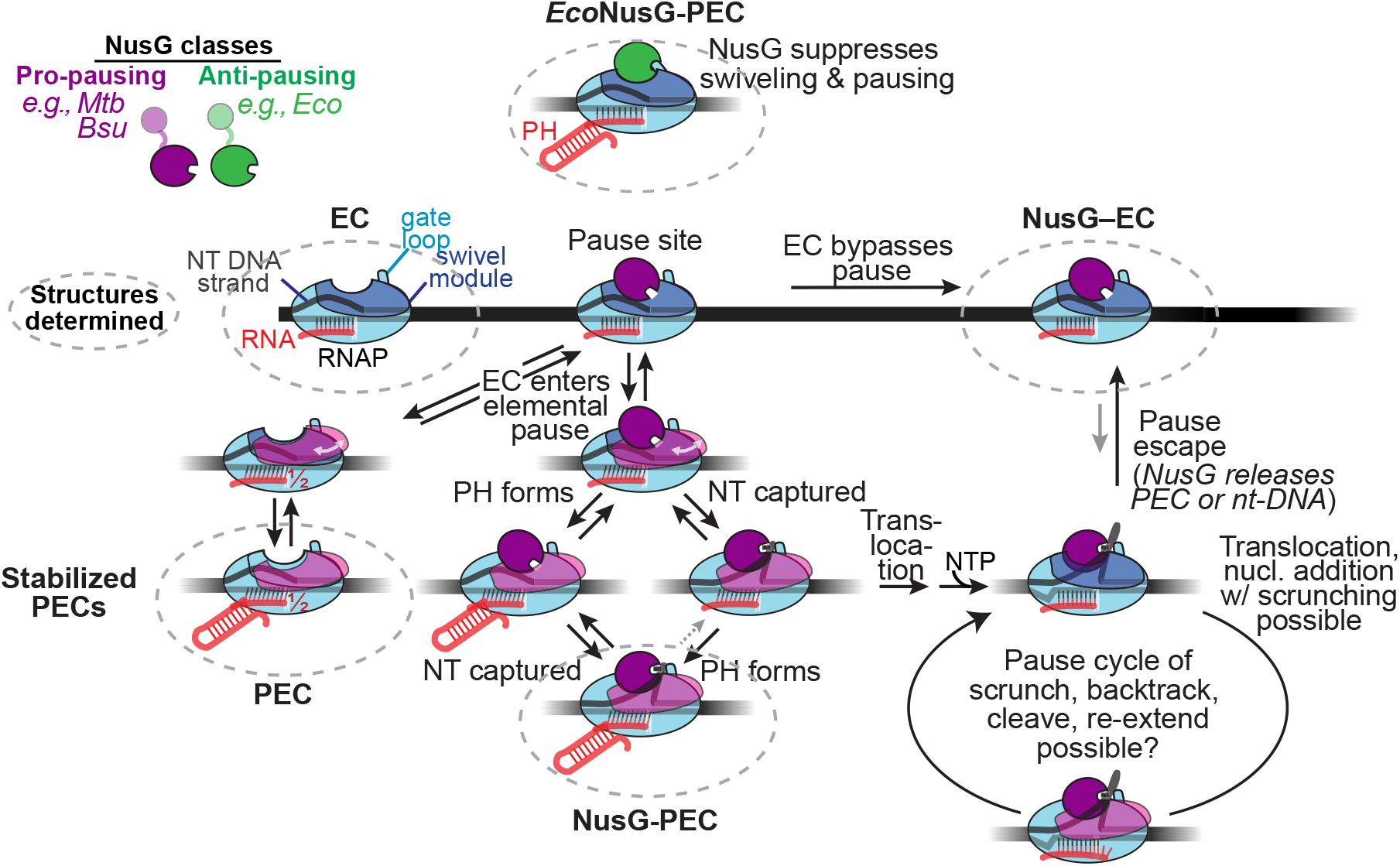
Effects of pro-pausing and anti-pausing NusG on transcription elongation. Intermediates whose cryo-EM structures are reported here are in dashed circles. Anti-pausing NusG suppresses pausing by inhibiting swiveling via an interaction with the GL. Pro-pausing NusG stimulates pausing by increasing the lifetime of swiveled states via interactions with the nt-DNA and RNAP protrusion. These interactions can be established either before or after the PEC is stabilized by formation of a PH. Once pro-pausing NusG stabilizes the PEC, it may remain in paused states during pausing cycles of limited RNA extension, DNA scrunching, backtracking, and transcript cleavage (Landick, 2021; Strobel and Roberts, 2015).

### Pro-pausing NusG may stimulate multiple steps in transcriptional pausing

Pausing is a multi-step mechanism that involves not only multiple conformations of the ePEC (Kang et al., 2022) but also multiple more long-lived states that are backtracked, stabilized by PHs or transcription factors, or even extended by one or more nucleotides by scrunching of the transcription bubble (Kang et al., 2019; Landick, 2021; Qian et al., 2021; Strobel and Roberts, 2015). The only universally conserved transcription factor, NusG/Spt5, can increase or decrease pausing through effects at more than one of these steps (Herbert et al., 2010; Kang et al., 2018b; Pasman and von Hippel, 2000; Wang and Artsimovitch, 2020; Yakhnin et al., 2020b). Our results reveal that pro-pausing *Mtb*NusG can act at the first steps of pausing, formation of the elemental and PH-stabilized PECs. *Mtb*NusG acts on *Mtb*RNAP as a pro-pausing factor, whereas *Eco*NusG acts on the same RNAP at the same DNA sites as an anti-pausing factor (Figure 1). *Mtb*NusG and *Eco*NusG accomplish these opposing effects by promoting or suppressing RNAP swiveling through differences in preferred interactions with the RNAP (Figures 3D and 3E).

Pro-pausing *Mtb*NusG stabilizes swiveling by: *i*) interacting with the RNAP βprotrusion, and *ii*) interacting with the single-stranded nt-DNA to promote the DNA wedge in the gap between the NusG-NGN and the βlobe (the gap itself created by swiveling; Figures 2B and 3D). The wedging of the DNA in this gap (promoted by *Mtb*NusG) further stabilizes swiveling (i.e. the DNA wedge blocks RNAP unswiveling in the presence of NusG).

Conversely, anti-pausing *Eco*NusG unswivels the PEC, overcoming the presence of the swivel-promoting PH by interacting with the βlobe/GL (instead of the βprotrusion; Figure 3E). Thus, the key to the striking functional difference between pro-pausing and anti-pausing NusG is the promotion or suppression (respectively) of swiveling, which appears to be mediated by NusG preference for interacting with the βprotrusion and the nt-DNA wedge (pro-pausing NusG) or for interacting with the βlobe–GL (anti-pausing NusG).

Pro-pausing NusG could also act at steps downstream of the initial pause by trapping RNAP in a pausing cycle (Figure 7) (Landick, 2021; Strobel and Roberts, 2015). Strong contacts to the nt-DNA could prevent translocation of the upstream DNA fork-junction. Scrunching of the DNA bubble could allow one or more rounds of nucleotide addition, as has been observed for Eco σ^70^ and the NusG paralog RfaH bound to PECs (Belogurov et al., 2010; Pukhrambam et al., 2022; Strobel and Roberts, 2015). During scrunching, the nt-DNA extrudes away from the ePEC. The gap between *Mtb*NusG and the protrusion could accommodate nt-DNA extrusion. The thermodynamic destabilization resulting from nucleotide addition and scrunching in the pause cycle would be relieved either by backtracking and transcript cleavage, as observed for σ^70^ and RfaH, or by escape from the pause cycle when NusG release of the nt-DNA or of RNAP altogether allows forward translocation (Figure 7).

It is currently unclear if the pro-pausing activities of *Mtb*NusG and the classically studied pro-pausing *Bsu*NusG are identical. In our assays, *Bsu*NusG exhibits stronger pro-pausing activity than *Mtb*NusG (Figures 1C and D). Further, their biological functions may differ. The existence of translationally coupled-attenuation mechanisms suggests transcription and translation are coupled in *Mtb* (D’Halluin et al., 2022; Lee et al., 2022) as they are in the model g-proteobacterium *Eco* where NusG mediates transcription–translation coupling (Burmann et al., 2010). *Bsu* transcription is uncoupled from translation (Johnson et al., 2020) and, unlike for *Mtb* (Figure 6), *Bsu*NusG is dispensable for growth. The TTnTTT motif, which is reported to be necessary for *Bsu*NusG pro-pausing activity, is not required for *Mtb*NusG pro-pausing activity (Figure 5D). However, the TTnTTT motif was defined using strong pauses that were not detected when NusG was absent (Yakhnin et al., 2020a). Our reanalysis of the same dataset focused instead on pauses that occur without NusG but are increased ~3-10-fold (i.e., pro-pausing NusG activity) reveals no sequence conservation in the region corresponding to the TTnTTT motif. It seems possible that the TTnTTT motif exhibits exceptional affinity for *Bsu*NusG that could increase strong pausing by trapping *Bsu*RNAP in pause cycles (Figure 7).

### Pro-pausing NusG activity may be crucial for robust growth of mycobacteria

The inducible CRISPRi and complementation system optimized in this study provides a platform for future studies of *nusG* in mycobacteria. Previous methods had limitations: the operon structure of the *nusG* locus complicates interpretation of genome-wide screen results, while NusG’s contributions to mycobacterial growth render it impossible to perturb endogenously (i.e., to generate knockouts). Here, we demonstrate the specific essentiality of *nusG* in mycobacteria for the first time and find that pro-pausing activity is crucial for robust mycobacterial growth. Our *in vivo* system should enable future genomics work to identify which loci are affected by this pro-pausing activity leading to the observed decrease in mycobacterial fitness.

## ACKNOWLEDGEMENTS

We thank members of the Darst-Campbell, Landick, and Rock Laboratories and N. Paknejad for helpful discussions; and M. Ebrahim, J. Sotiris, and H. Ng at The Rockefeller University, and K. Maruthi and K. Neselu at NYSBC Evelyn Gruss Lipper Cryo-electron Microscopy Resource Center for help with cryo-EM data collection. Some of the work was performed at the Simons Electron Microscopy Center and National Resource for Automated Molecular Microscopy, located at the New York Structural Biology Center, supported by grants from the Simons Foundation (SF349247), New York State Office of Science, Technology and Academic Research, and the NIH National Institute of General Medical Sciences (GM103310). This work was supported by NIH grants R35 GM118130 to S.A.D., NIH/NIAID New Innovator Award 1DP2AI144850-01 to J.M.R., R01 GM114450 to E.A.C., and R01 GM38660 to R.L.

## Author contributions

Conceptualization: M.D., E.O.O., R.F., J.R., E.A.C., R.L.; Transcription assays and protein purification: E.O.O., M.D., R.A.M; Recombinant DNA generation: M.D., E.O.O., R.F., S.K., R. A.M.; Cryo-EM specimen preparation: M.D.; Cryo-EM data collection: M.D.; Cryo-EM data processing: M.D., R.F., M.L.; Model building and structural analysis: M.D., R.F., M.L., J.J.B., S. A.D., E.O.O, R.L., E.A.C.; *In vivo* studies in mycobacteria: R.F., S.K., J.R., M.D., E.A.C.; Funding acquisition and supervision: J.R., S.A.D, E.A.C., R.L.; Manuscript first draft: M.D., E.O.O., R.F., S.A.D., E.A.C., R.L. All authors contributed to finalizing the written manuscript.

## Resource Availability

### Lead contact

More information and requests for resources and reagents should be directed to the Lead contact, Robert Landick.

### Materials availability

All unique materials generated in this study are available by contacting the Lead contact.

## Resource Availability

### Materials availability

All unique materials generated in this study are available by contacting the Lead contact.

### Data and code availability

The cryo-EM density maps have been deposited in the Electron Microscopy Data Bank under accession codes EMD-27956 (EC), EMD-27935 (PEC*-Mtb*NusG), EMD-27938 (PEC*-Eco*NusG), EMD-27944 (PEC) and EMD-27942 (EC*-Mtb*NusG). The atomic coordinates have been deposited in the Protein Data Bank under accession codes 8E95 (EC), 8E74 (PEC*-Mtb*NusG), 8E79 (PEC*-Eco*NusG), 8E8M (PEC), and 8E82 (EC*-Mtb*NusG). These datasets are publicly available as of the date of publication. Any additional information required to reanalyze the data reported in this paper is available from the lead contact upon request.

## Experimental model and subject details

### Mycobacterium smegmatis (Msm)

All *Msm* strains were derived from strain mc^2^155, a strain that has lost the function of the *eptC* gene, thereby leading to highly efficient plasmid transformation (Panas et al., 2014). *Msm* were grown at 37°C in Difco Middlebrook 7H9 broth (BD #271310) or 7H10 (BD #262710) plates supplemented with 0.2% glycerol (7H9) or 0.5% glycerol (7H10), 0.05% Tween80, and 1X albumin dextrose catalase (ADC). Where required, antibacterial or small molecules were used at the following concentrations: kanamycin (KAN) at 20 μg ml^−1^; nourseothricin (NAT) at 25 μg ml^−1^; and anhydrotetracycline (ATc) at 100 ng ml^−1^. Small (5mL) liquid cultures were grown standing in tissue culture flasks, while larger (50mL) cultures were grown shaking in Erlenmeyer flasks at 120 rpm.

## Methods details

Structural biology software was accessed through the SBGrid consortium (Morin et al., 2013).

### Protein expression and purification

#### *Mtb* NusG

Plasmid pAC82 was used to overexpress WT *Mtb* NusG (Czyz et al., 2014). pAC82 is kanamycin-resistant, contains the T7 promoter, ten histidine residues, and a precision protease cleavage site upstream of NusG. Plasmids encoding NusG mutants were generated using Q5 Site-directed mutagenesis (NEB) and sequenced to confirm the presence of target mutations. Plasmids encoding different versions of *Mtb* NusG were grown in *Eco* Rosetta2 (Novagen) cells in LB with 50 μg kanamycin/mL at 37 °C to an OD_600_ of 0.4, then transferred to room temperature and left shaking to an OD_600_ of 0.6. Protein expression was induced by adding IPTG to a final concentration of 0.5 mM, grown for additional 6 hr, then harvested by centrifugation (8,000 x *g*, 15 min at 4 °C). Harvested cells were resuspended in 20 mM Tris-HCl, pH 8.0, 500 mM NaCl, 0.1 mM PMSF, 1 mM protease inhibitor cocktail (Roche), 0.3 mg lysozyme/mL, 5 mM β-mercaptoethanol, 5% glycerol and lysed by sonication. The lysate was centrifuged (27,000 x *g*, 15 min, 4 °C) and the supernatant incubated with batch Ni^2+^-NTA agarose resin (GoldBio) while tumbling for 1 h at 4 °C. The mixture was poured into a disposable column (Qiagen) and the flow-through collected using gravity filtration. The column was washed with 20 CV of the wash buffer (50 mM Tris-HCl, pH 8.0, 500 mM NaCl, 10 mM imidazole, 5% glycerol) and NusG eluted from the column using the elution buffer (20 mM Tris-HCl, pH 8.0, 500 mM NaCl, 200 mM Imidazole, 5% glycerol). The eluted protein was subsequently purified by gel filtration chromatography on a HiLoad Superdex 26/600 200 pg (GE Healthcare Life Sciences) in 20 mM Tris-HCl, pH 8.0, 200 mM NaCl, 1 mM EDTA, 1 mM DTT, 5% glycerol, 0.1% Triton X-100. Eluted protein was incubated with human rhinovirus (HRV) 3C protease at 4 °C for 4 h to cleave the N-terminal His_10_-tag and then passed over Ni^2+^-NTA agarose to remove uncleaved protein. Additional glycerol was added to the purified NusG to a final concentration of 25% (v/v), and the sample aliquoted and flash frozen in liquid nitrogen. Aliquots were stored in –80 °C until use.

#### *Mtb* RNAP

*Mtb* RNAP was purified as previously described (Lilic et al., 2021). Plasmid pMP61 was used to overexpress WT *Mtb* RNAP. pMP61 is kanamycin-resistant, contains a T7 promoter, followed by *rpoA, rpoZ*, a linked *rpoBC* and a His_8_-tag. *rpoB* and *rpoC* were fused using a short linker to minimize the possibility of *Eco* β or β’ intermixing with *Mtb* subunits. pMP61 was grown in *Eco* Rosetta2 cells in LB with 50 μg kanamycin/mL and 34 μg chloramphenicol/mL at 37 °C to an OD_600_ of 0.3, then transferred to room temperature and left shaking to an apparent OD_600_ of 0.6. RNAP expression was induced by adding IPTG to a final concentration of 0.1 mM, grown overnight for 16 hr, and harvested by centrifugation (8000 g, 15 min at 4 °C). Harvested cells were resuspended in 50 mM Tris-HCl, pH 8.0, 1 mM EDTA, 1 mM PMSF, 1 mM protease inhibitor cocktail, 5% glycerol and lysed by sonication. The lysate was centrifuged (27,000 x *g*, 15 min, 4 °C) and polyethyleneimine (PEI, Sigma-Aldrich) added to the supernatant to a final concentration of 0.6% (w/v) and stirred for 10 min to precipitate DNA binding proteins including target RNAP. After centrifugation (11000x*g*, 15 min, 4 °C), the pellet was resuspended in PEI wash buffer (10 mM Tris-HCl, pH 7.9, 5% v/v glycerol, 0.1 mM EDTA, 5 μM ZnCl2, 300 mM NaCl) to remove non-target proteins. The mixture was centrifuged (11000g, 15 min, 4 °C), supernatant discarded, then RNAP eluted from the pellet into PEI Elution Buffer (10 mM Tris-HCl, pH 7.9, 5% v/v glycerol, 0.1 mM EDTA, 5 μM ZnCl2, 1 M NaCl). After centrifugation, RNAP was precipitated from the supernatant by adding (NH4)2SO4 to a final concentration of 0.35 g/L. The pellet was dissolved in Nickel buffer A (20 mM Tris pH 8.0, 5% glycerol, 1 M NaCl, 10mM imidazole) and loaded onto a HisTrap FF 5 mL column (GE Healthcare Life Sciences). The column was washed with Nickel buffer A then RNAP eluted with Nickel elution buffer (20 mM Tris, pH 8.0, 5% glycerol, 1 M NaCl, 250 mM imidazole). Eluted RNAP was subsequently purified by gel filtration chromatography on a HiLoad Superdex 26/600 200 pg in 10mM Tris pH 8.0, 5% glycerol, 0.1mM EDTA, 500mM NaCl, 5mM DTT. Eluted samples were aliquoted, flash frozen in liquid nitrogen and stored in −80 °C until usage.

#### *Mtb* σA–RbpA

*Mtb* σA–RbpA was purified as previously described with slight modifications (Czyz et al., 2014; Hubin et al., 2017). The *Mtb* σA expression vector is derived from the pCDFDuet-1 backbone (Novagen) and contains the T7 promoter, twelve histidine residues, and a precision protease cleavage site upstream of *Mtb* σA. The *Mtb* RbpA vector is derived from the pet21C backbone (Novagen) and contains the T7 promoter upstream of untagged *Mtb* RbpA. Both plasmids were co-transformed into *Eco* BL21(DE3) cells (Novagen). Protein expression was induced by adding IPTG to a final concentration of 0.5 mM when cells reached an apparent OD_600_ of 0.6, followed by overnight growth at 18°C, then harvested by centrifugation (4,000 g, 15 min at 4 °C). Harvested cells were resuspended in 50 mM Tris-HCl, pH 8.0, 500 mM NaCl, 5mM imidazole, 0.1 mM PMSF, 1 mM protease inhibitor cocktail, and 1 mM β-mercaptoethanol, then lysed using a continuous-flow French press. The lysate was centrifuged twice (16,000g, 30 min, 4 °C) and the proteins were purified by Ni^2+^-affinity chromatography (HisTrap IMAC HP, GE Healthcare Life Sciences, Pittsburgh, PA) via elution at 50 mM Tris-HCl, pH 8.0, 500 mM NaCl, 500mM imidazole, and 1 mM β-mercaptoethanol. Following elution, the complex was dialyzed overnight into 50mM Tris-HCl, pH 8.0, 500mM NaCl, 5mM imidazole, and 1mM β-mercaptoethanol and the His_12_ tag was cleaved with precision protease overnight at a ratio of 1/30 (protease mass/cleavage target mass). The cleaved complex was loaded onto a second Ni^2+^-affinity column and was retrieved from the flow-through. The complex was loaded directly onto a size exclusion column (SuperDex-200 16/16, GE Healthcare Life Sciences) equilibrated with 50mM Tris-HCl, pH 8, 500mM NaCl, and 1mM DTT. The sample was concentrated to 4mg/mL by centrifugal filtration and stored at −80 °C until usage.

#### *Mtb* CarD

*Mtb* CarD was purified as previously described (Boyaci et al., 2019; Srivastava et al., 2013). In brief, *Mtb* CarD was amplified from genomic DNA and cloned into the overexpression vector pET SUMO (Invitrogen) and transformed into *Eco* BL21(DE3) cells (Novagen). Protein expression was induced by adding IPTG to a final concentration of 1 mM when cells reached an apparent OD_600_ of 0.6, followed by 4 hours of growth at 28°C, then harvested by centrifugation (4,000 g, 15 min at 4 °C). Harvested cells were resuspended in 20 mM Tris-HCl, pH 8.0, 150 mM K-Glutamate, 5mM MgCl2, 0.1 mM PMSF, 1 mM protease inhibitor cocktail, and 1 mM b-mercaptoethanol, then lysed using a continuous-flow French press. The lysate was centrifuged twice (16,000g, 30 min, 4 °C) and the proteins were purified by Ni^2+^-affinity chromatography (HisTrap IMAC HP, GE Healthcare Life Sciences, Pittsburgh, PA) via elution at 20 mM Tris-HCl, pH 8.0, 150mM K-Glutamate, 250mM imidazole, and 1 mM β-mercaptoethanol. Following elution, the complex was dialyzed overnight into 20mM Tris-HCl, pH 8.0, 150mM K-Glutamate, 5mM MgCl2, and 1mM β-mercaptoethanol and the His_10_ tag was cleaved with ULP-1 protease (Invitrogen) overnight at a ratio of 1/30 (protease mass/cleavage target mass). The cleaved complex was loaded onto a second Ni^2+^-affinity column and was retrieved from the flowthrough. The complex was loaded directly onto a size exclusion column (SuperDex-200 16/16, GE Healthcare Life Sciences) equilibrated with 20mM Tris-HCl, pH 8, 150mM K-Glutamate, 5mM MgCl2 and 2.5mM DTT. The sample was concentrated to 5mg/mL by centrifugal filtration and stored at −80°C.

#### *Bsu* NusG

Plasmid pAY94 was used to overexpress WT *Bsu* NusG. pAY94 plasmid was a generous gift from Paul Babitzke (Yakhnin et al., 2008). pAY94 is Ampicillin-resistant, contains the T7 promoter, and six histidine residues upstream of NusG. *Bsu* NusG was purified using the same protocol described above for *Mtb* NusG except the purified *Bsu* NusG was not subjected to HRV 3C protease to cleave off His_6_-tag.

#### *Eco* NusG

*Eco* NusG was purified as described previously (Mooney et al., 2009). Briefly, plasmid pRM1160, an overexpression plasmid for *Eco* NusG was transformed and grown in *Eco* BLR (DE3) cells in LB with 100 μg/mL ampicillin at 37° C to an OD_600_ of 0.35 – 0.5, induced for protein expression by addition of IPTG to a final concentration of 10 mM, grown for an additional 2.5 – 3 hours, harvested by centrifugation. Harvested cells were resuspended at 10 mL/g in lysis buffer (20 mM Tris-HCl, pH 7.8, 3 mM EDTA, 5 mM β-mercaptoethanol, 100 mM NaCl, 20 mg/mL PMSF), lysozyme was added to 0.2 mg/mL, and the cells were incubated for 10 min at 22 °C, and then for 20 min on ice. Unless stated otherwise, all subsequent steps were performed at 4 °C. Sodium deoxycholate was added to a final concentration of 0.06% (w/v), and the resulting solution incubated for 20 min, then NaCl was added to 0.3 M. The lysate was sonicated 4 times for 15 s at 22 °C, with 3 min on ice between sonication treatments. Polymin P was added to a final concentration of 0.6%, and the lysate incubated for 20 min and centrifuged at 12000g. Ammonium sulfate was added to the supernatant at a final concentration of 50% and stirred for 30 min. The lysate was centrifuged for 20 min at 12,000 x *g*, and the pellet resuspended in 30 mL of buffer Q (10 mM Tris-HCl, pH 7.8, 1 mM EDTA, 1 mM DTT, 5% glycerol). The lysate was then dialyzed against 4 × 1 L changes of buffer Q for a total of 16 h and spun at 12000g, and the supernatant was loaded onto a HisTrap FF 5 mL column (GE Healthcare Life Sciences). The column was washed with buffer Q then NusG eluted with elution buffer (10 mM Tris-HCl, pH 7.8, 1 mM EDTA, 5% glycerol, 250 mM imidazole). Eluted NusG was subsequently purified by Q ion-exchange chromatography, aliquoted, flash frozen in liquid nitrogen and stored in −80 °C until usage.

#### Promoter-based *In vitro* transcription assay

The DNA sequence used in the promoter-based assays was PCR amplified from *Bsu* chromosomal DNA and cloned into pMiniT (NEB) to generate pRM1244 plasmid. This sequence is from the untranslated leader of *Bsu* trp operon and harbors the U108 pause and the U105 terminated positions described herein. The template for *in vitro* transcription was obtained by PCR amplification of pRM1244 plasmid such that it only had the consensus –10 element, extended –10 element and not the –35 element. This template also contained a C-less cassette (+1 to +29), U105 terminated and U108 pause positions. Core RNAP was incubated with σA/RbpA for 10 min at 37 °C to form holo-RNAP, followed by 10 min incubation with CarD at 37 °C. To initiate transcription, holo-RNAP (50 nM) was incubated with template DNA (10 nM), ApU dinucleotide (150 μM), ATP + UTP (both at 2.5 μM), and 1 μM [α-32P]GTP (130 Ci/mmol) in transcription buffer (10 mM Hepes, pH 8.0, 50 mM KGlutamate, 10 mM MgOAc, 0.1 mM EDTA, 5 μg/mL Acetylated BSA and 1 mM DTT) for 15 min at 37 °C to form a halted complex at G29. Different NusGs were added at this step at a final concentration of 1 μM and transcription restarted by adding a master mix containing NTP mix (A + C + G + U), Heparin and KGlutamate at a final concentration of 150 μM, 50 μg/mL and 100 mM respectively. Aliquots were removed at respective timepoints, and transcription stopped using a 2X Stop buffer (8 M urea, 50 mM EDTA, 90 mM Tris-Borate buffer, pH 8.3, 0.02% bromophenol blue, 0.02% xylene cyanol). Samples were then analyzed on an 8% denaturing PAGE (19:1 acrylamide: bis acrylamide, 45 mM Tris-borate, pH 8.3, 1.25 mM Na2EDTA, 8M urea) for 2 h at 50 W, the gel exposed on a Storage Phosphor Screen and imaged using a Typhoon PhosphoImager (GE Healthcare).

#### Scaffold-based *In vitro* transcription assay

PEC RNA was 5’ ^32^P-labeled by treatment with [g-^32^P] ATP polynucleotide kinase. T-DNA (1.33 μM) and the labelled RNA (1 μM) were pre-assembled in transcription buffer and incubated on a thermocycler for 2 min at 95 °C. The mixture was then subsequently cooled down to 75 °C for 2 min, 45 °C for 5 min and from 43 – 23 °C for 2 min per lowering 1 °C. After incubation, the mixture was either used immediately or stored in the −20 °C freezer. PECs were reconstituted at the U42 pause site by incubation of 100 nM RNA42, 133 nM t-DNA, and 200 nM *Mtb* core RNAP in 30 μL buffer for 5 min at 37 °C. Nt-DNA (400 nM) was added, and the mixture incubated for an additional 5 min at 37 °C. Heparin (50 μg/mL) was added to the mixture to prevent non-specific interactions between core RNAP and nucleic acids. After incubation with 1 μM NusG, RNA42 was extended by addition of ATP + UTP + 3’-dGTP (All at a final concentration of 100 μM), and aliquots (5 μL) removed after 10, 20, 30, 60 and 120 sec and combined with 5 μL 2X stop buffer (8 M urea, 50 mM EDTA, 90 mM Tris-Borate buffer, pH 8.3, 0.02% bromophenol blue, 0.02% xylene cyanol). The remaining fraction was chased by addition of ATP + UTP + 3’-dGTP to a final concentration of 600 μM and reaction stopped as described for the different timepoints. Samples were then analyzed on a 15% denaturing PAGE (19:1 acrylamide: bis acrylamide, 45 mM Tris-borate, pH 8.3, 1.25 mM Na2EDTA, 8M urea) for 4 - 5 h at 50 W, the gel exposed on a Storage Phosphor Screen and imaged using a Typhoon PhosphoImager (GE Healthcare).

#### *Msm* plasmid construction and cloning

plRL61 (Addgene, #163633) was used to generate *Msm* CRISPRi *nusG* knockdown strain, as previously described by previously (Bosch et al., 2021; Rock et al., 2017). plRL61 integration into the *Msm* chromosome is mediated by the L5 integrase, supplied in trans on a separate nonreplicating suicide vector, plRL19 (Addgene, #163634). The CRISPRi plasmid backbone was digested using BsmBI–v2 (New England Biolabs, #R0739L) and agarose gel–purified. A single-guide RNA (sgRNA) bearing appropriate overhangs and designed to hybridize to the nontemplate strand of the *nusG* open reading frame was annealed and ligated (T4 ligase, New England Biolabs, #M0202M) into the digested backbone. Identification of desired clones was confirmed by Sanger sequencing.

The CRISPRi *nusG* knockdown strain was prepared by electroporation of plasmids into *Msm*. A 20-ml culture of *Msm* was grown to an apparent OD_600_ of 0.8 to 1.0 and pelleted (4000*g*, 10 min) before being washed three times with 10% glycerol (20 ml) and resuspended in 1 ml of 10% glycerol. For each transformation, 50 μl of electrocompetent cells was combined with ~200-500 ng of CRISPRi plasmid DNA and ~100ng plRL19 before electroporation in a 2-mm cuvette (Bio-Rad, #1652082) using the Gene Pulser X cell electroporation system (Bio-Rad, #1652660) set at 2500 V, 700 ohms, and 25 μF. Electroporated cells were recovered in 7H9 broth for at least 2 hours and 20 minutes before being plated on 7H10 agar supplemented with KAN (20 μg/ml) for 3 days to select for transformants prior to being imaged.

Complementation constructs were generated in a Tweety-integrating (plRL91) backbone. All constructs contain the genomic region between coordinates 1437738 and 1440768, based on *Msm* reference genome assembly NC008596.1. This start coordinate was chosen to include the promoter *“pnusG* operon”, which was selected based on conservation and high expression levels between two independent *Msm* TSS datasets (Li et al., 2017; Martini et al., 2019). The end coordinate was chosen to include the *rplK* and *rplA* loci, which we predicted may be cotranscribed with *nusG*, whereas the subsequent gene downstream of *rplA* is in the opposite orientation. All “test” *nusG* alleles were inserted into this complementation construct backbone using Gibson assembly. Specifically, the CRISPRi-resistant complementation allele was amplified from *Msm* genomic DNA using Gibson assembly primers encoding synonymous mutations in the complementation allele protospacer adjacent motif and sgRNA seed sequence to prevent sgRNA targeting (d’Andrea et al., 2022). Homolog NGN boundaries were chosen by generating fusion constructs and assessing their likely secondary and tertiary structures using ColabFold (Mirdita et al., 2022). The final fusion constructs contained the following amino acids: *Msm* N-terminal extension = 1-89; *Mtb* NGN = 49-152; *Bsu* NGN = 2-115; *Eco* NGN = 2-120; and *Msm* linker and KOW = 201-280. Homolog NGN fragments were amplified from genomic DNA of the relevant species. The amplified fragments were then cloned into DraIII-HF (New England Biolabs, #R3510S)–digested plRL91. Identification of desired clones was confirmed by whole-plasmid sequencing with Plasmidsaurus. plRL91-derived constructs were co-transformed into *Msm* with plRL62, a nonreplicating vector supplying the Tweety integrase in trans. All plasmids used for *Msm* experiments are listed in supplemental data file 3.

#### Immunoblotting

*Msm* cultures were growth-synchronized before ATc was added to a final concentration of 100 ng ml^−1^. For each time point, at least 10 OD_600_ units of culture were harvested by centrifugation (4000*g*, 10 min) and resuspended in 600 μl of ice-cold lysis buffer [50 mM tris and 50 mM NaCl (pH 7.4)] before lysis by bead-beating in Lysis B Matrix tubes (MP Biomedicals, #116911050) using a Precellys Evolution homogenizer (Bertin Instruments, #P000062-PEVO0-A; 3 × 10,000 rpm, 30-s intervals). The cell lysates were cleared by centrifugation (*20,000g*, >10 min), and a 20-μl aliquot was incubated with 4× lithium dodecyl sulfate (LDS) sample buffer (Invitrogen, #NP0007) supplemented with 100 mM dithiothreitol at 70°C for 10 min. Samples were separated on a 4 to 12% bis-tris polyacrylamide gel (Invitrogen, #NP0323BOX) in MES running buffer, transferred to a nitrocellulose membrane using the TransBlot Turbo Transfer System (Bio-Rad, #1704150), and incubated for 1 hour in blocking buffer (LI-COR, #927-60001). Lysates with 3xFLAG-tagged proteins were probed with anti-RpoB (BioLegend, #663905) or anti-FLAG (Sigma-Aldrich, #F3165) primary antibodies overnight at 4°C and subsequently detected with fluorescent goat anti-mouse secondary antibody (Bio-Rad, #12004159). Endogenous *Msm* NusG was probed with polyclonal antibodies raised against *Eco* NusG overnight at 4°C and subsequently detected with fluorescent goat anti-rabbit secondary antibody (Bio-Rad, #12004162).

#### Nucleic acid sequences used for cryo-EM samples

The scaffold-based sequences used for cryo-EM are listed below. DNA oligos were ordered gel purified from Integrated DNA Technologies, Coralville, IA. Protected RNA oligos were ordered from Dharmacon, Inc.

**PEC**:

nt-DNA

5’ CGTCAGAAAGAAAACCCTTTATTTGTTATATAGTATTTTATCCTCTCATGCCGG 3’

t-DNA

5’ CCGGCATGAGAGGATAAAATACTATATCCTGGTTAAAGGGTTTTCTTTCTGACG 3’

RNA

5’ AUUCUACCCAAAAGAAGUCUUUCUUUUGGGUUUAACCAGGAU 3’

**EC:**

nt-DNA

5’ CGTCGGATACTTGCGGGCTAGCCTCTTTTGGCGGCGAATACCCTCTCATGCCGG 3’

t-DNA

5’ CCGGCATGAGAGGGTATTCGCCGCCTACCTCTCCTAGCCCGCAAGTATCCGACG 3’

RNA:

5’ GCAUUCAAAGCGGAGAGGUA 3’

#### Cryo-EM grid preparation

C-flat holey carbon grids (CF-1.2/1.3-4Au, Photochips, Morrisville, NC) were glow-discharged for 20 s before the application of 3.5 μL of the samples. OG, (n-Octyl-b-D-Glucoside, Anatrace, Maumee, OH), was added to the samples to a final 0.1% (w/v). Using a Vitrobot Mark IV (Thermo Fisher Scientific Electron Microscopy, Hillsboro, OR), grids were blotted and plunge frozen into liquid ethane with 100% chamber humidity at 22 °C.

#### Preparation of ECs and PECs for cryo-EM

Synthetic DNA oligonucleotides were obtained from Integrated DNA Technologies (Coralville, IA), RNA oligonucleotides from GE Healthcare Dharmacon Inc. (Lafayette, CO). The nucleic acids for the EC and PEC scaffolds were resuspended in annealing buffer (10 mM HEPES pH 7.5, 1 mM EDTA, 50 mM NaCl). Respective template DNA (t-DNA) and RNA were annealed in a 1:3 ratio for 5 min at 95°C in a covered heat block, then cooled to room temperature, then immediately moved to ice. The annealed RNA-DNA hybrids were stored at −80°C until use. Purified *Mtb* core RNAP was dialyzed overnight into cryo-EM buffer (20 mM HEPES pH 7.5, 150 mM KGlu, 5 mM MgOAc, 2.5 mM DTT). To prepare complexes, annealed t-DNA:RNA was combined with core at 1.3:1 molar excess and incubated at room temperature for 15 min. Next, non-template DNA (nt-DNA) was added to the complexes at 1.5:1 molar excess with the t-DNA:RNA and incubated for 10 min at room temperature. The complexes were then injected onto the Superose 6 Increase 10/300 GL column (Cytiva) for purification. *Mtb* NusG or *Eco* NusG was buffer exchanged and purified on the Superose 6 column. Purified NusG was added in 3x molar excess to purified core RNAP and incubated for 10 min at room temperature. Complexes were concentrated by centrifugal filtration (Amicon Ultra) to 4-6.5 mg RNAP/mL before grid preparation.

#### Cryo-EM data acquisition and processing

Grids were imaged using a 300 keV Titan Krios (Thermo Fisher Scientific Electron Microscopy) equipped with a K2 Summit direct electron detector (Gatan, Pleasanton, CA) or K3 Gatan direct electron detector. Dose-fractionated movies were recorded in counting mode using Leginon (Nicholson et al., 2010). Pixel size, defocus range, dose rate, and total exposure values for each data collection are found in Table S1. Dose-fractionated movies were gain-normalized, drift-collected, summed, and dose-weighted using MotionCor2 (Zheng et al., 2017). The CTF (Contrast Transfer Function) was estimated for individual summed images using Patch CTF estimation in cryoSPARC2/3 (Punjani et al., 2017). Blob picker in was used in cryoSPARC2/3 (Punjani et al., 2017) to pick particles with no input template, and particles were extracted in cryoSPARC2/3 using a box size of 300. For the number of extracted particles in each dataset, please reference the individual pipelines in Figures S2A, S2E, S3A, S3E, and S4A. Ab initio models were obtained and used as a 3D template. cryoSPARC2/3 adaption of “random phase 3D classification” (Gong et al., 2016) heterogeneous refinements were performed to curate 3D model and classify “junk” particles. Curated particles from each dataset were refined using Local CTF refinement (Punjani et al., 2017) and polished in RELION (Scheres, 2012; Zivanov et al., 2018). Heterogeneous refinements were performed on polished particles in cryoSPARC2/3 to separate unwanted classes. Note: depending on the dataset, the last heterogeneous refinement step was performed either before or after polishing (indicated for each dataset in Figures S2-S4). Final classifications were further refined using cryosparc2/3 non-uniform refinement. Number of particles and nominal resolutions for each of the final maps are found in Figures S2A, S2E, S3A, S3E, and S4A (in dashed boxes). Nominal resolutions were determined by using the gold-standard FSC 0.143 cutoff (Afonine et al., 2018).

#### Model building, refinement, and validation

For initial models of the complexes, the *Mtb* RNAP cryo-EM structure (PDB ID: 6EDT (Boyaci et al., 2019) were manually fit into the cryo-EM density maps using UCSF Chimera (Pettersen et al., 2004). For real space refinement, rigid body refinement followed by all-atom refinements with Ramachandran and secondary structure restraints. Refined models were analyzed and modified in Coot (Emsley and Cowtan, 2004). Remote 3DFSC Processing Server was used for processing Fourier shell correlations of cryo-EM maps (Tan et al., 2017).

#### Contacts analysis

*Mtb* and *Eco* NusG surface contact areas with RNAP were analyzed using PISA (Protein interfaces, surfaces, and assemblies’) service at the European Bioinformatics Institute. (http://www.ebi.ac.uk/pdbe/prot_int/pistart.html) (Krissinel and Henrick, 2007). The source code used to identify amino-acid contacts is available online at Github (https://github.com/darst-campbell-lab/residue-contacts-app).

## Quantification and Statistical Analysis

### Promoter-based pause quantification

Synthesized RNA bands were quantified using Image J software (NIH). The first step was to draw straight vertical lines for each lane that start a few inches below the U105 and U108 sites and extend to include the run-off RNA products. A graphical representation of the drawn line enabled the isolation of specific RNA bands, herein, divided into: U105, U108 or run-off bands. The total intensity was calculated by adding these three intensities. Termination at U105 and pausing at U108 were represented as percentages of the total intensity, which were compared over time (figure 1). The fold effect of different NusGs was determined by taking the intensity of target band in the presence of a test NusG and dividing by the intensity when no NusG is present.

### Scaffold-based pause quantification

Synthesized RNA bands were quantified using Image J software (NIH). For all timepoints, the RNA intensity for paused and escaped RNAP were measured separately and the shorter transcripts resulting from RNA degradation were not included during analysis. The paused RNA intensity of the chased sample was subtracted from each timepoint to correct for inactive complexes. The corrected pause RNA was plotted as a function of time and the plot used to determine the pausing kinetics by fitting to a single-exponential decay function (figure 1)

### Quantification used for cryo-EM

For calculations of Fourier shell correlations (FSC) in Figures S2C, S2G, S3C, S3G, and S4C, the FSC cut-off criterion of 0.143 (Afonine et al., 2018) was used. Quantifications and statistical analyses for model refinement and validation were calculated in PHENIX (Afonine et al., 2018).

## Key Resource Table

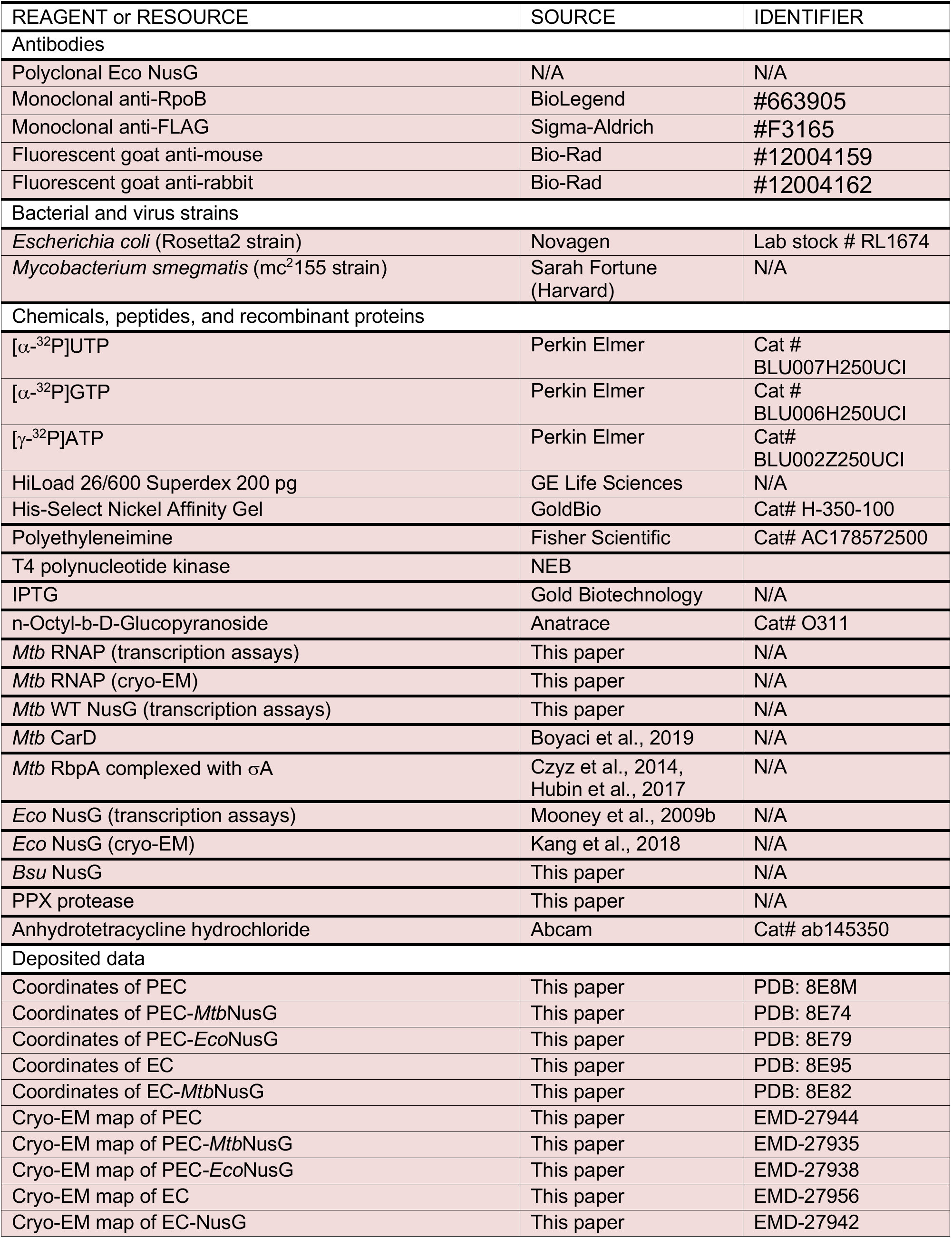
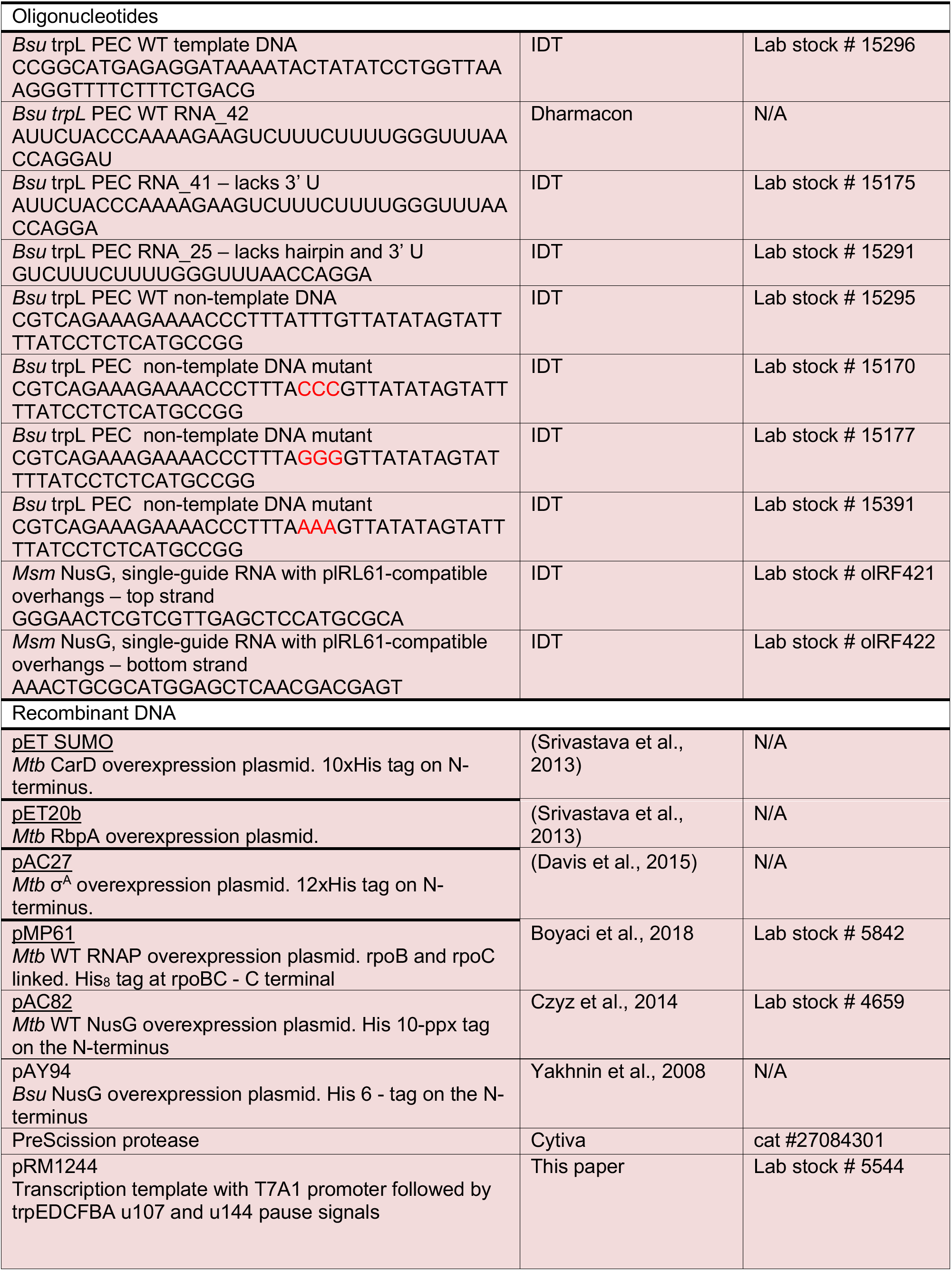
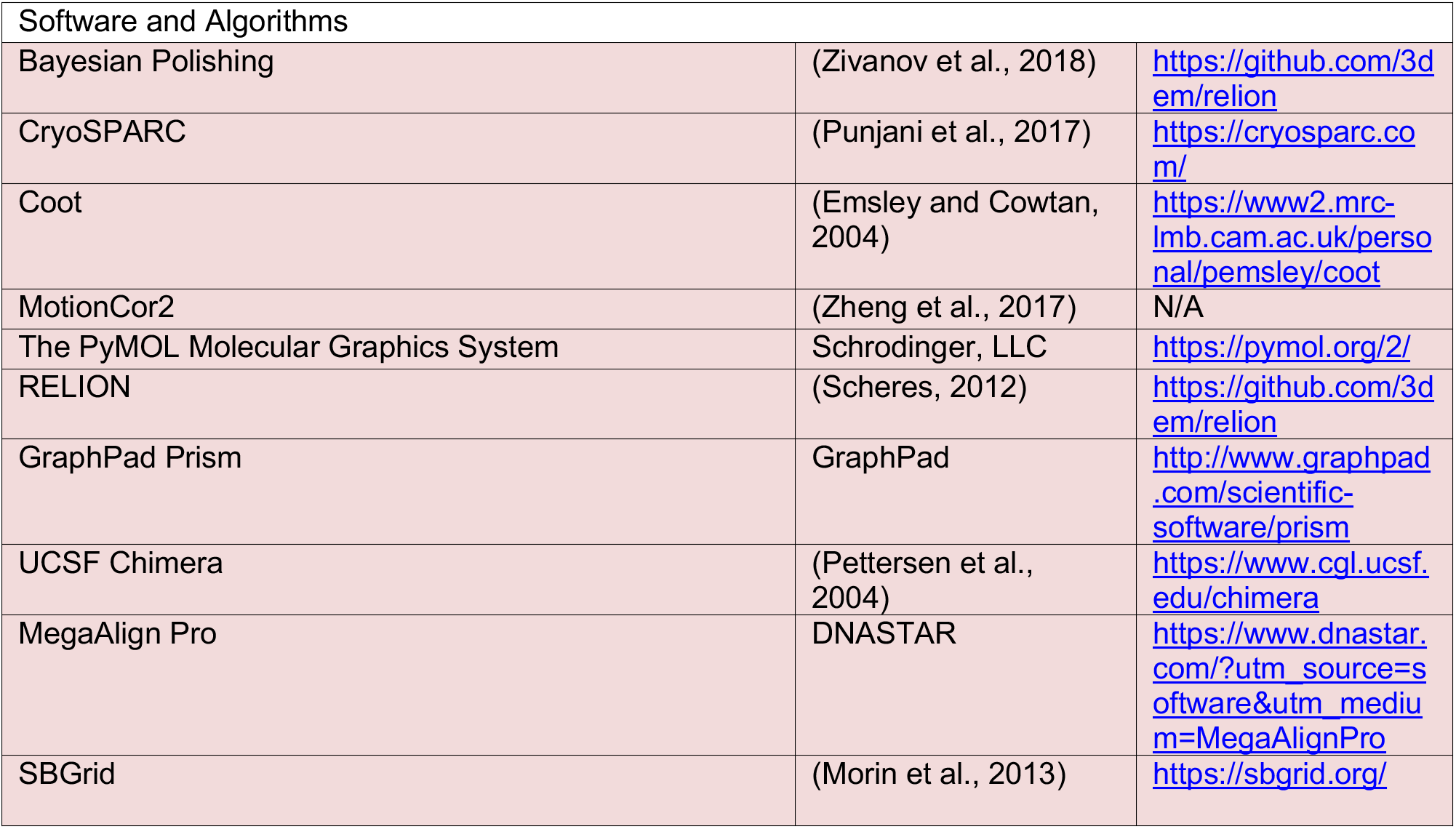

