## Supplemental Figures and Table for "Structural and functional basis of the universal transcription factor NusG pro-pausing activity in *Mycobacterium tuberculosis*"

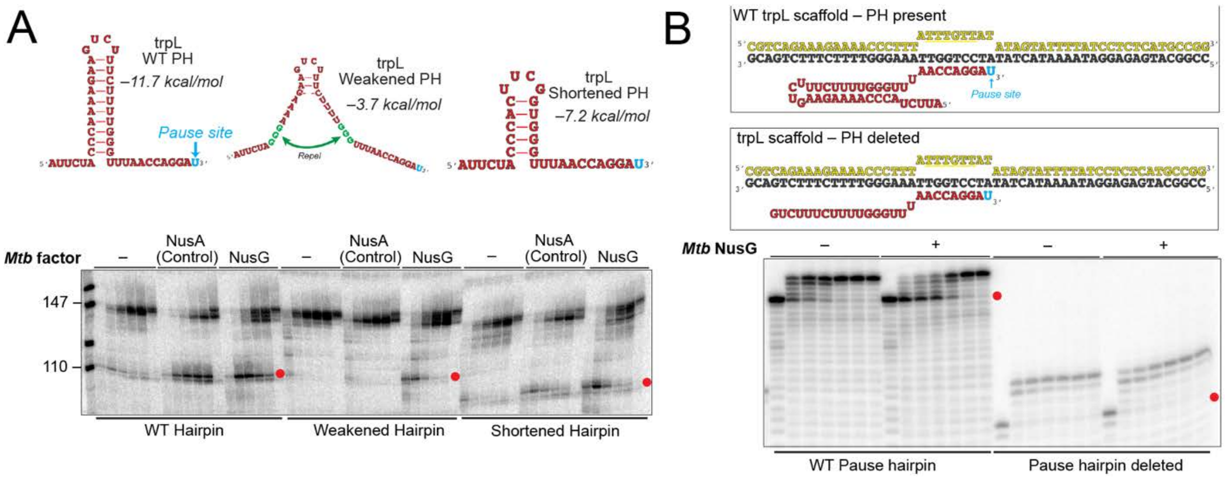


**Figure S1. Mtb NusG enhances pausing of RNAP at the hairpin-stabilized trpL pause.** (A) Schematic of the synthesized RNA sequence from the promoter-based assay showing a pause hairpin upstream of the U108 pause site (left). The pause hairpin is weakened by disrupting the tri-C-G base pairs at the bottom of the stem (middle) and shortened by deleting five base pairs at the top of the stem (right*). Mtb*NusG strongly enhances pausing of RNAP at U108 when the WT pause hairpin is present, but this enhancement is reduced when the pause hairpin is weakened or shortened. NusA was used as control to confirm that the pause hairpin was successfully disrupted. PH stands for pause hairpin. Timepoints collected were 1, 2, 4, 10 and 15 mins. The last lane is the chased sample. (B) WT scaffold used to obtain PEC cryoEM structures (top) and its variation that lacks the pause hairpin (bottom). *Mtb*NusG enhances pausing when the WT pause hairpin is present, but this enhancement is completely lost when the pause hairpin is deleted. Timepoints collected were 0, 10, 20, 30, 60, and 120s. Last lane is the chased sample.

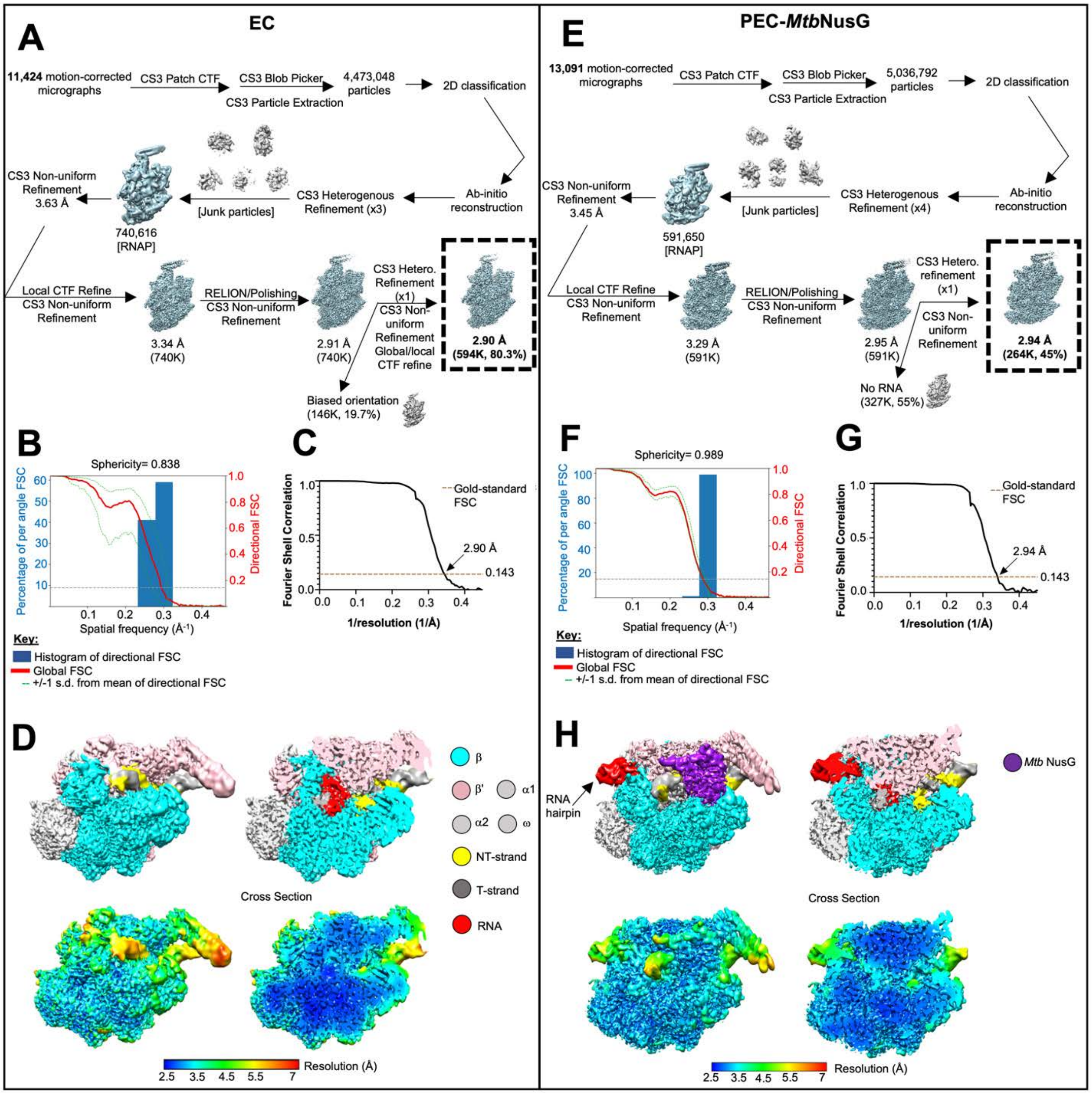


**Figure S2. Data processing workflow and cryo-EM map validation for EC and PEC-*Mtb*NusG**

**(A)** Cryo-EM pipeline used for processing EC. Dose-fractionated movies (11,424) were frame aligned and summed using MotionCor2 (Zheng et al., 2017). Motion-corrected micrographs were processed in cryoSPARC3 (CS3) (Punjani et al., 2017). CTFs were estimated using Patch CTF, and the Blob Picker was used to pick particles, which were subsequently extracted. Extracted particles were curated using 3 rounds of CS3 heterogeneous refinement (6 classes each) using an adaption of “random-phase 3D classification” (Gong et al., 2016). CS3 local CTF refinement was then performed on curated particles, and the particles were polished in RELION (Scheres, 2012; Zivanov et al., 2018). Polished particles were refined with CS3 non-uniform refinement. Refined particles were then classified using CS3 heterogeneous refinement to remove unwanted particles (biased orientations).

**(B)** Histogram and directional FSC plot for EC was calculated using the Remote 3DFSC Processing Server (Tan et al., 2017). Global FSC is shown with a red line.

(**C)** Gold-standard FSC was calculated from CS3. The dashed line represents FSC cutoff of 0.143, which indicates a nominal resolution of 2.90 Å.

(**D)** The EC cryo-EM density map is colored according to the key (Pettersen et al., 2004). The right view is a cross-section of the left view. The bottom maps show local resolution calculations from CS3.

**(E)** Cryo-EM pipeline used for processing PEC-*Mtb*NusG. Dose fractionated movies (13,091) were frame aligned and summed using MotionCor2 (Zheng et al., 2017). Motion-corrected micrographs were processed in cryoSPARC3 (CS3) (Punjani et al., 2017). CTFs were estimated using Patch CTF, and the Blob Picker was used to pick particles, which were subsequently extracted. Extracted particles were curated using 4 rounds of CS3 heterogeneous refinement (6 classes each) using an adaption of “random-phase 3D classification” (Gong et al., 2016). CS3 local CTF refinement was then performed on curated particles, and the particles were polished in RELION (Scheres, 2012; Zivanov et al., 2018). Polished particles were refined with CS3 non-uniform refinement. Refined particles were then classified using CS3 heterogeneous refinement to remove unwanted particles (those without RNA).

**(F)** Histogram and directional FSC plot for EC was calculated using the Remote 3DFSC Processing Server (Tan et al., 2017). Global FSC is shown with a red line.

**(G)** Gold-standard FSC was calculated from CS3. The dashed line represents FSC cutoff of 0.143, which indicates a nominal resolution of 2.94 Å.

**(H)** The PEC-*Mtb*NusG cryo-EM density map is colored according to the key (same as Figure S2D) (Pettersen et al., 2004). The right view is a cross-section of the left view. The bottom maps show local resolution calculations from CS3.

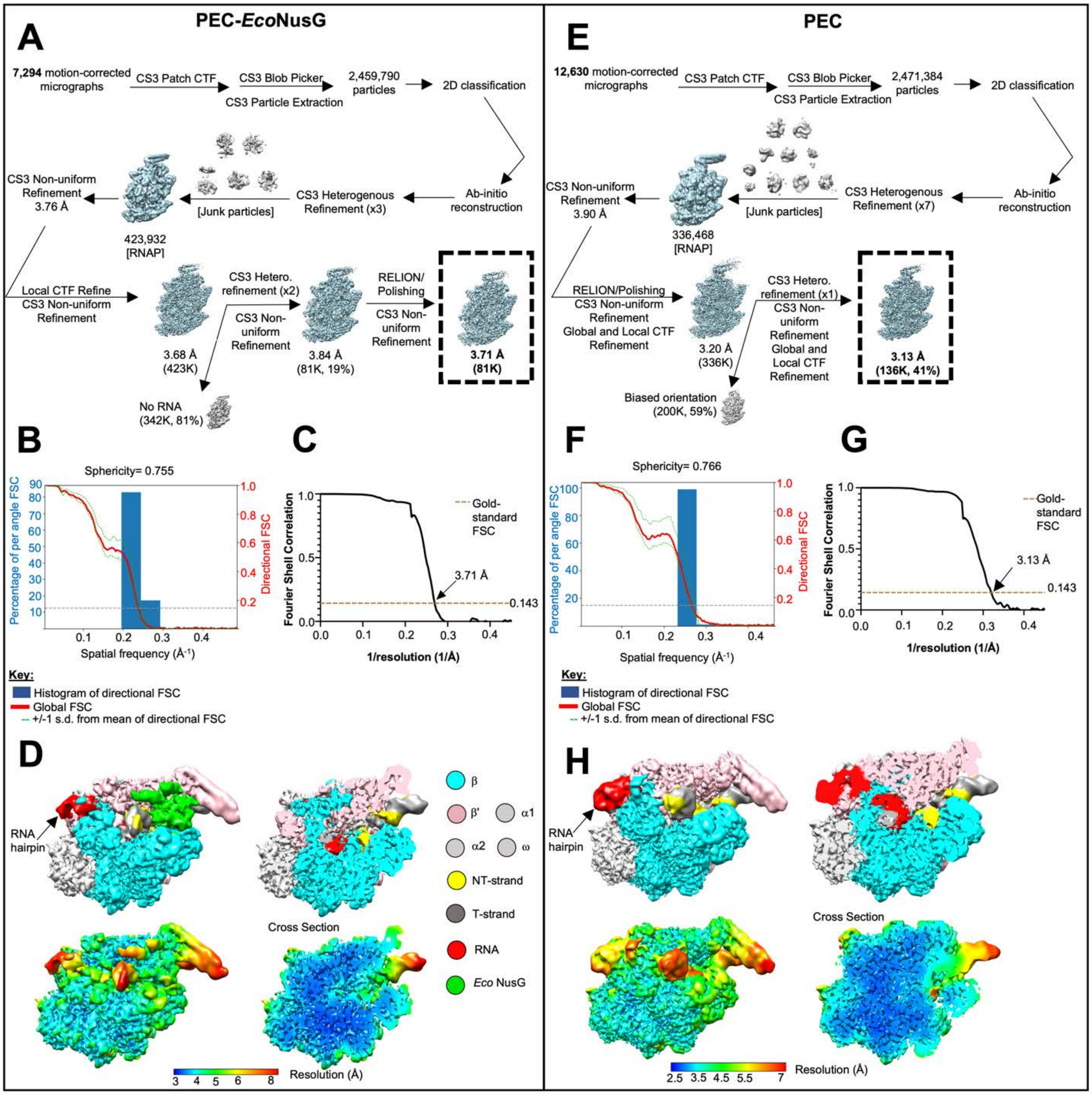


**Figure S3. Data processing workflow and cryo-EM map validation for PEC-*Eco*NusG and PEC**

**(A)** Cryo-EM pipeline used for processing PEC-*Eco*NusG. Dose-fractionated movies (7,294) were frame aligned and summed using MotionCor2 (Zheng et al., 2017). Motion-corrected micrographs were processed in cryoSPARC3 (CS3) (Punjani et al., 2017). CTFs were estimated using Patch CTF, and the Blob Picker was used to pick particles, which were subsequently extracted. Extracted particles were curated using 3 rounds of CS3 heterogeneous refinement (6 classes each) using an adaption of "random-phase 3D classification" (Gong et al., 2016). CS3 local CTF refinement was then performed on curated particles. The particles were then sorted with CS3 heterogeneous refinement to remove unwanted particles lacking RNA. The remaining particles were polished in RELION (Scheres, 2012; Zivanov et al., 2018) and refined with CS3 non-uniform refinement.

**(B)** Histogram and directional FSC plot were calculated using the Remote 3DFSC Processing Server (Tan et al., 2017). Global FSC is shown with a red line.

(**C)** Gold-standard FSC was calculated from CS3. The dashed line represents FSC cutoff of 0.143, which indicates a nominal resolution of 3.71 Å.

**(D)** The PEC-*Eco*NusG cryo-EM density map is colored according to the key (Pettersen et al., 2004). The right view is a cross-section of the left view. The bottom maps show local resolution calculations from CS3.

**(E)** The cryo-EM pipeline used for processing PEC. Dose-fractionated movies (12,630) were frame aligned and summed using MotionCor2 (Zheng et al., 2017). Motion-corrected micrographs were processed in cryoSPARC3 (CS3) (Punjani et al., 2017). CTFs were estimated using Patch CTF, and the Blob Picker was used to pick particles, which were subsequently extracted. Extracted particles were curated using 7 rounds of CS3 heterogeneous refinement (12 classes each) using an adaption of “random-phase 3D classification” (Gong et al., 2016). Curated particles were then polished in RELION (Scheres, 2012; Zivanov et al., 2018) and refined with CS3 non-uniform refinement (including global and local CTF refinements). Polished/refined particles were refined further with CS3 heterogeneous refinement to remove unwanted particles (biased orientations).

**(F)** Histogram and directional FSC plot for PEC was calculated using the Remote 3DFSC Processing Server (Tan et al., 2017). Global FSC is shown with a red line. **(G)** Gold-standard FSC was calculated from CS3. The dashed line represents FSC cutoff of 0.143, which indicates a nominal resolution of 3.13 Å.

**(H)** The PEC cryo-EM density map is colored according to the key (same as Figure S3D) (Pettersen et al., 2004). The right view is a cross-section of the left view. The bottom maps show local resolution calculations from CS3.

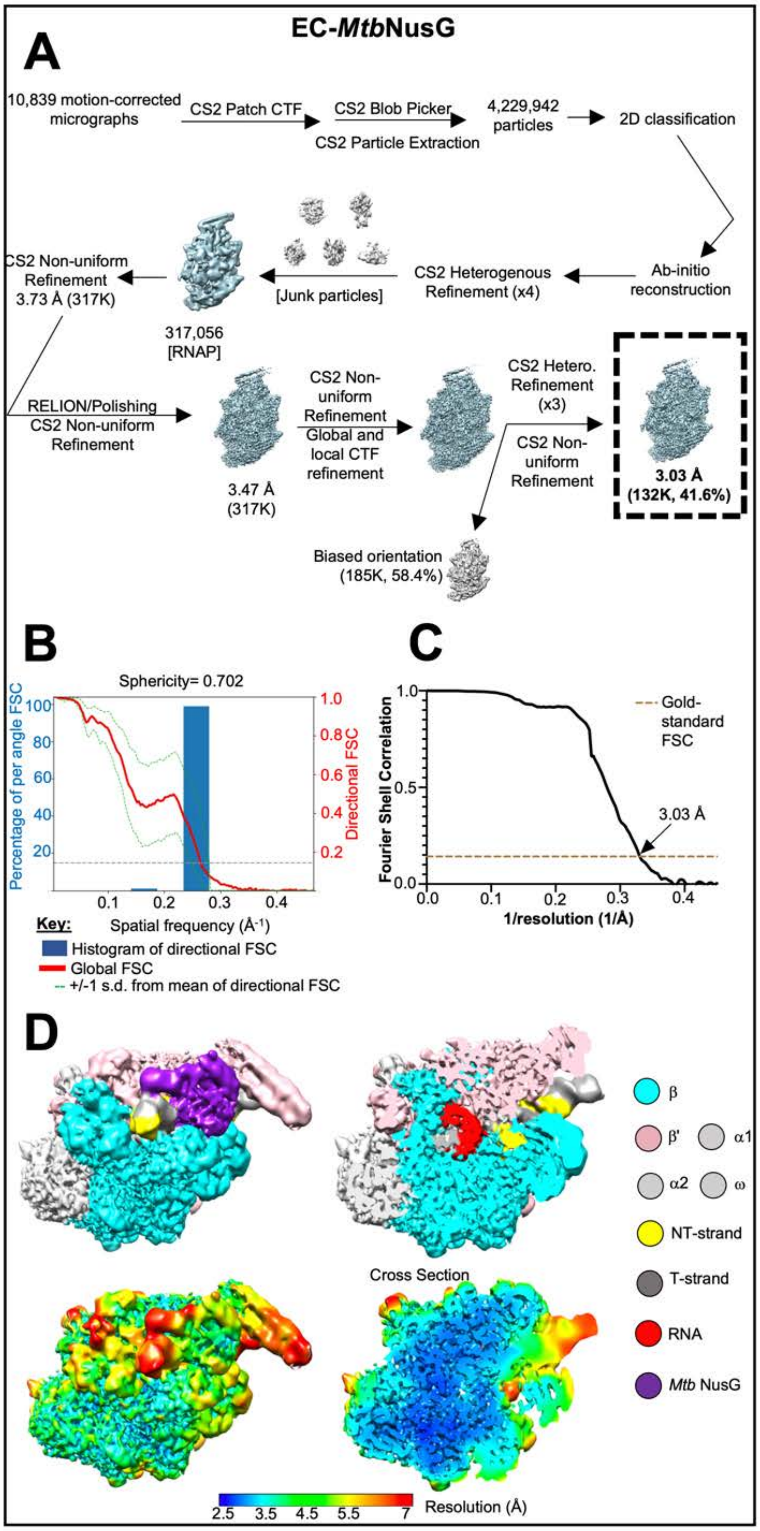


**Figure S4. Data processing workflow and cryo-EM map validation for EC-*Mtb*NusG**

**(A)** Cryo-EM pipeline used for processing EC-*Mtb*NusG. Dose-fractionated movies (10,839) were frame aligned and summed using MotionCor2 (Zheng et al., 2017). Motion-corrected micrographs were processed in cryoSPARC2 (CS2) (Punjani et al., 2017). CTFs were estimated using Patch CTF, and the Blob Picker was used to pick particles, which were subsequently extracted. Extracted particles were curated using 4 rounds of CS2 heterogeneous refinement (6 classes each) using an adaption of “random-phase 3D classification” (Gong et al., 2016). Curated particles were polished in RELION (Scheres, 2012; Zivanov et al., 2018). Polished particles were refined with CS2 non-uniform refinement (with global and local CTF refinements). Refined particles were then classified using CS2 heterogeneous refinement to remove unwanted particles (biased orientations).

**(B)** Histogram and directional FSC plot for EC-*Mtb*NusG was calculated using the Remote 3DFSC Processing Server (Tan et al., 2017). Global FSC is shown with a red line.

(**C)** Gold-standard FSC was calculated from CS2. The dashed line represents FSC cutoff of 0.143, which indicates a nominal resolution of 3.03 Å.

**(D)** The EC-*Mtb*NusG cryo-EM density map is colored according to the key (Pettersen et al., 2004). The right view is a cross-section of the left view. The bottom maps show local resolution calculations from CS2.

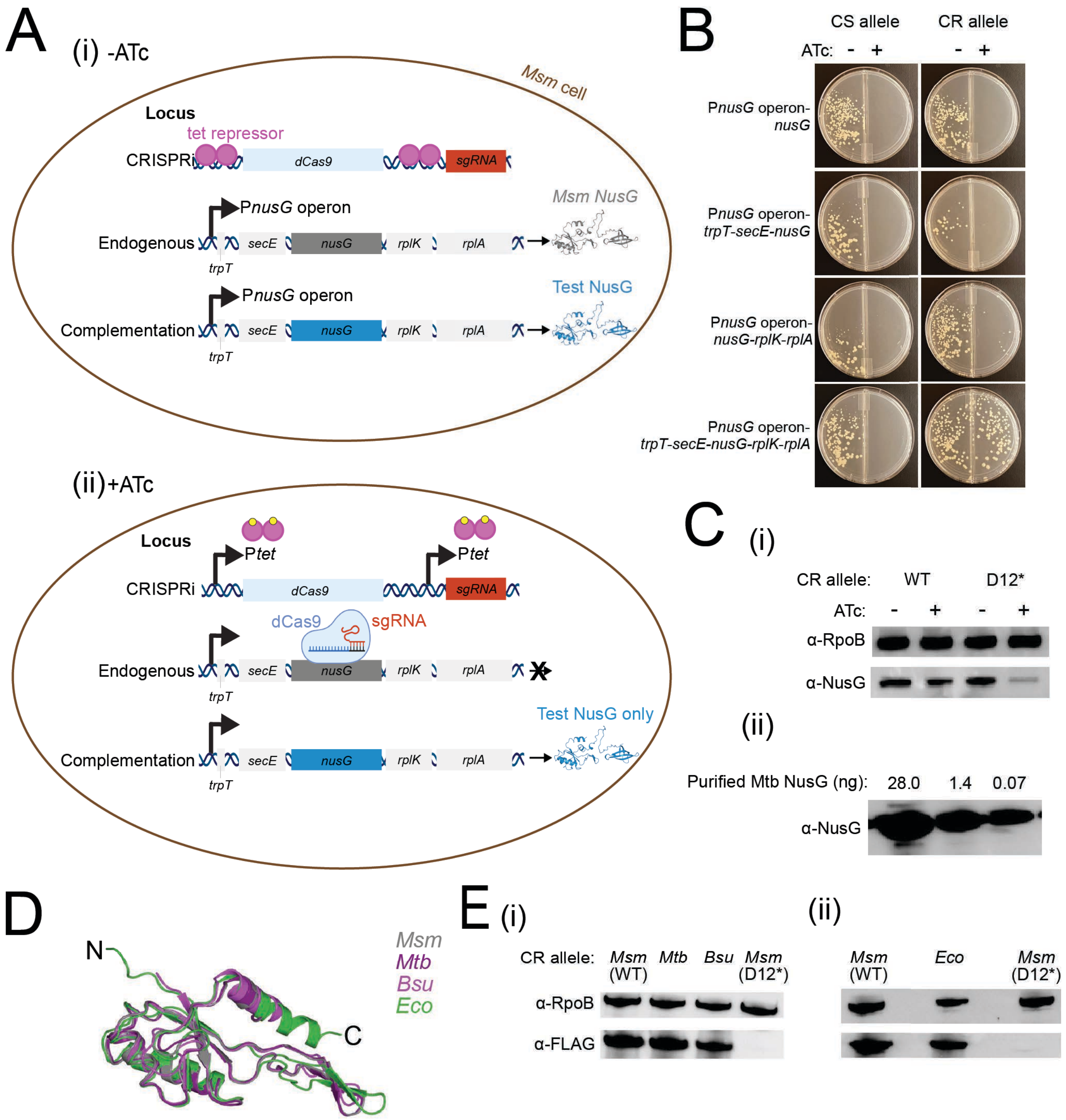


**Figure S5. Design and quality control of *in vivo* systems to test the effects of NusG on *M. smegmatis* (Msm) fitness.**
(**A**) *nusG* CRISPRi knockdown and complementation system schematic. DNA and dCas9 schematics were generated in BioRender. NusG protein images represent a ColabFold structure prediction of *Msm*NusG, with disordered amino acids 1-74 removed for simplicity.
(**B**) Five-day outgrowth of CRISPRi-sensitive (CS) or CRISPRi-resistant (CR) complementation constructs transformed into the *nusG* operon knockdown strain.
I (i) Western blot of NusG levels in Msm after 12 hours of ATc exposure, run on the same gel as (ii) dilutions of purified *Mtb*NusG protein.
(**D**) ColabFold structural predictions and breakpoints for NusG NGNs that were fused to the *Msm* N-terminal extension and *Msm* linker to KOW domains for use in complementation assays. Only the NGN is shown from the ColabFold prediction for simplicity.

(**E**) Western blots of N-terminally tagged 3xFLAG-GSGG NusG constructs. (i) Western blot of lysates harvested from liquid culture +ATc after 18 hours. (ii) Western blot of lysates scraped from agar plates in the absence of ATc after three days (*Msm* WT and D12*) or six days (*Eco*). *Eco* plates were harvested later to obtain equivalent biomass.

**Table S1. Cryo-EM data collection, refinement, and validation statistics**

| | **EC** PDB ID: 8E95 | **PEC-*Mtb*NusG** PDB ID: 8E74 | **PEC-*Eco*NusG** PDB ID: 8E79 | **PEC** PDB ID: 8E8M | **EC-*Mtb*NusG** PDB ID: 8E82 |
|---|---|---|---|---|---|
| **Data collection and processing** | | | | | |
| Magnification | 81,000 | 81,000 | 29,000 | 81,000 | 64,000 |
| Voltage (kV) | 300 | 300 | 300 | 300 | 300 |
| Electron exposure (e-/Å$^2$) | 51.24 | 63.89 | 51.84 | 51.14 | 64.78 |
| Defocus range (mm) | 0.8 to 1.8 | 1.0 to 2.0 | -1.5 to -2.5 | 1.0 to 2.5 | 1.0 to 2.5 |
| Pixel size (Å) | 1.0825 | 1.0825 | 1.03 | 1.0825 | 1.076 |
| Symmetry imposed | C1 | C1 | C1 | C1 | C1 |
| Initial particle images (no.) | 4,473,048 | 5,036,792 | 2,459,790 | 2,471,384 | 4,229,942 |
| Final particle images (no.) | 593,911 | 264,224 | 80,855 | 136,516 | 132,324 |
| Map resolution (Å) FSC threshold 0.143 | 2.90 | 2.94 | 3.71 | 3.13 | 3.03 |
| Map resolution range (Å) | 2.5-7 | 2.5-7 | 3-8 | 2.5-7 | 2.5-7 |
| **Refinement** | | | | | |
| Initial model used (PDB code) | 6EDT | 6EDT | 6EDT | 6EDT | 6EDT |
| Model resolution range (Å) | 2.5-7 | 2.5-7 | 3-8 | 2.5-7 | 2.5-7 |
| Map sharpening B factor (A$^2$) | 96.8 | 108.0 | 128.0 | 105.4 | 70.0 |
| **Model composition** | | | | | |
| Non-hydrogen atoms | 47,342 | 50,726 | 50,322 | 48,355 | 49,255 |
| Protein residues | 2,909 | 3,048 | 3,017 | 2,924 | 3,031 |
| Nucleic acid residues | 68 | 106 | 107 | 93 | 68 |
| Ligands | 3 (1 $Mg^{2+}$, 2 $Zn^{2+}$) | 3 (1 $Mg^{2+}$, 2 $Zn^{2+}$) | 3 (1 $Mg^{2+}$, 2 $Zn^{2+}$) | 3 (1 $Mg^{2+}$, 2 $Zn^{2+}$) | 3 (1 $Mg^{2+}$, 2 $Zn^{2+}$) |
| **B factors (A$^2$)** | | | | | |
| Protein | 30.72 | 58.54 | 96.03 | 75.35 | 77.98 |
| Nucleic acid | 80.14 | 134.8 | 180.87 | 174.03 | 139.82 |
| Ligands | 58.97 | 90.52 | 118.48 | 105.99 | 119.53 |
| **R.m.s. deviations** | | | | | |
| Bond lengths (Å) | 0.003 | 0.004 | 0.004 | 0.004 | 0.003 |
| Bond angles (°) | 0.568 | 0.644 | 0.622 | 0.562 | 0.635 |
| **Validation** | | | | | |
| MolProbity score | 2.13 | 1.91 | 1.83 | 1.85 | 2.28 |
| Clashscore | 4.67 | 4.71 | 7.00 | 4.47 | 7.27 |
| Poor rotamers (%) | 4.35 | 2.59 | 3.79 | 2.37 | 3.89 |
| **Ramachandran plot[a]** | | | | | |
| Favored (%) | 95.06 | 94.85 | 93.13 | 95.12 | 93.40 |
| Allowed (%) | 4.67 | 4.82 | 6.70 | 4.60 | 6.21 |
| Outliers (%) | 0.28 | 0.33 | 0.17 | 0.27 | 0.40 |

[a] Ramachandran plot parameters from PHENIX
